# Sister chromatid cohesion establishment during DNA replication termination

**DOI:** 10.1101/2022.09.15.508094

**Authors:** George Cameron, Dominika Gruszka, Sherry Xie, Kim A Nasmyth, Madhusudhan Srinivasan, Hasan Yardimci

## Abstract

The cohesin complex tethers sister chromatids together from the moment they are generated in S-phase until their separation in anaphase^1,2^. This fundamental phenomenon, called sister chromatid cohesion, underpins orderly chromosome segregation. The replisome complex coordinates cohesion establishment with replication of parental DNA^3^. Cohesion can be established by cohesin complexes bound to DNA before replication^4,5^, but how replisome interaction with pre-loaded cohesin complexes results in cohesion is not known. Prevailing models suggest cohesion is established by replisome passage through the cohesin ring or by transfer of cohesin behind the replication fork by replisome components^5^. Unexpectedly, by visualising single replication forks colliding with pre-loaded cohesin complexes, we find that cohesin is pushed by the replisome to where a converging replisome is met. Whilst the converging replisomes are removed during DNA replication termination, cohesin remains on nascent DNA. We demonstrate that these cohesin molecules tether the newly replicated sister DNAs together. Our results support a new model where sister chromatid cohesion is established during DNA replication termination, providing important insight into the molecular mechanism of cohesion establishment.

## Introduction

Cohesin is a ring-shaped structural maintenance of chromosomes (SMC) complex, with four core subunits (SMC1, SMC3, RAD21^Scc1^ and SA1/SA2^Scc3^)^6^. In addition to the essential role of cohesin in sister chromatid cohesion, cohesin organises interphase chromosomes by loop extrusion^7,8^. Cohesin is loaded onto chromatin by the NIPBL/MAU2 (Scc2/4) loader^9^, which in vertebrates interacts with pre-replication complexes (pre-RCs)^10,11^. In eukaryotes, origins of replication are licensed during G1-phase through the formation of pre-RCs, which contain inactive double hexamers of MCM2-7^12^. When S-phase begins, pre-RCs are remodelled to form CDC45-MCM2-7-GINS (CMG) helicases, which then unwind DNA. The replisome complex, containing further components required for replication of chromatin, is assembled around the CMG helicase^13,14^. Once loaded onto DNA, the cohesin ring can interact topologically and non-topologically with DNA^15^. The observation that cleavage of RAD21 by separase in anaphase is essential for chromosome disjunction led to the notion that sister DNAs may be topologically trapped within cohesin rings^16^. This idea is further supported by analysis of sister minichromosome DNAs trapped inside covalently closed cohesin rings in budding yeast, where there is a perfect correlation between cohesion and co-entrapment of sister DNAs within cohesin rings^17^. Cohesion establishment is thought to occur only during S-phase^18^ and several replisome-associated proteins are important for cohesion establishment^4^. Cohesion establishment is therefore functionally coupled to DNA replication. Importantly, it is generated by two independent pathways ^4,19^. A ‘conversion’ pathway uses pre-loaded cohesin complexes associated with unreplicated (parental) DNA ahead of the replication forks to form cohesive structures, while a ‘*de novo*’ pathway uses cohesin rings loaded onto DNAs during S phase^14^. The molecular mechanisms by which cohesion is generated by these two pathways are unclear.

Two types of mechanism have been envisaged for conversion. Cohesion could be generated by passage of the replisome through cohesin rings that had previously entrapped unreplicated DNA (Supplementary Fig. 1a, left-hand side). Alternatively, cohesin rings could be transferred from unreplicated to replicated DNAs in a process requiring transient removal and ring re-opening while remaining associated with the replisome, as occurs for parental histones^20^ (Supplementary Fig. 1a, right-hand side). A prediction of both these scenarios is that DNA-associated cohesin rings, upon encounter with the replisome, would get incorporated into the replicated DNA and remain behind advancing replication forks. Testing this prediction *in vivo* has been challenging because cohesin is constantly mobile on DNA due to its loop extrusion activity and transcription mediated relocalisation^21^. Further, new cohesin is loaded onto DNA during most of the cell cycle^22^, making it difficult to follow pre-loaded cohesin complexes in cells. Finally, the stochasticity of eukaryotic DNA replication in individual cells leads to difficulty in assessing the outcome of cohesin-replisome encounters in a population of cells^12^. To overcome these limitations and determine the fate of pre-loaded cohesin during DNA replication, we performed live visualisation of replication forks encountering cohesin complexes at the single-molecule level.

## Results

### The replisome pushes pre-loaded cohesin during DNA replication

To study the outcome of replisome collision with cohesin in a physiologically relevant setting, we used *Xenopus laevis* egg extracts^23–25^, which contain all factors needed for *in vitro* DNA replication and repair^26^ whilst supporting cohesin loading and cohesion establishment^2^. Two different extracts allow a single round of DNA replication to be performed. High speed supernatant (HSS) is used to license DNA with pre-RCs in a sequence-independent manner, while nucleoplasmic extract (NPE) is used to replicate DNA from pre-RCs. A single-molecule assay to visualise replication of surface-immobilised DNA in egg extracts was previously developed, where total internal reflection fluorescence (TIRF) microscopy is used to visualise fluorescent molecules on stretched λ DNAs^24,25,27^. We used this assay to assess the outcomes of replication fork encounters with cohesin (Supplementary Fig. 1b). To visualise DNA-bound cohesin, we fluorescently labelled recombinant *Xenopus laevis* cohesin through a Halo tag on SMC3 (Supplementary Fig. 1c). Labelled cohesin was loaded onto chromatin in extracts in a manner dependent on DNA being licensed with MCM complexes, similar to endogenous cohesin loading^10,11^ (Supplementary Fig. 1d). Cohesin labelled with Janelia Fluor 646 (JF646-cohesin) was loaded onto surface-tethered λ DNAs either in licensing (HSS) extracts or in buffer supplemented with the cohesin loader (NIPBL-C) and ATP. DNA replication was initiated with NPE and replication fork progression followed by observing nascent DNA with fluorescently tagged Fen1 (Fen1-mKikGR), an enzyme involved in lagging-strand processing. To monitor encounter of individual replication forks with pre-loaded cohesin, origin firing was restricted using p27^kip^, a CDK inhibitor (Supplementary Fig. 1b).

Prevailing models for cohesin conversion predict incorporation of pre-loaded cohesin into nascent DNA behind the replication fork. Unexpectedly, in our conditions, this outcome of cohesin transfer was observed only in a small minority of events (5% of the events). (Supplementary Figs. 2a,b and 3a-c). In most cases, we instead observed that most commonly pre-loaded cohesin was pushed ahead of replication forks (cohesin sliding, 66% of events). In 20% of events, cohesin was removed shortly after encounter with the replication fork and in 9% of events replication fork stalling was observed upon encounter with pre-loaded cohesin. The lack of cohesin transfer was surprising as extracts contain all factors needed for cohesion establishment by the conversion pathway.

We next investigated if endogenous cohesin in *Xenopus* extracts interferes with the transfer of DNA-loaded fluorescently-tagged cohesin during replication. Previously we have shown that parental histone transfer behind replication forks is reduced by the high concentrations of soluble histones present in *Xenopus* extracts^28^. Free histones likely inhibit parental histone interaction with replisome components and prevent efficient histone transfer, raising the possibility endogenous cohesin in *Xenopus* extracts could have a similar inhibitory effect on cohesin transfer. To test this, endogenous cohesin was immunodepleted from extracts used for replication, which had little or no effect on fork speeds observed during replication (Supplementary Figs. 3d and 4a). Importantly, cohesin depletion did not elevate the rate of cohesin transfer and most cohesin was still pushed by forks (Supplementary Figs. 3c and 4b-d). This result suggests that endogenous cohesin complexes in extracts do not prevent pre-loaded cohesin from being transferred by replisomes.

We also considered if the presence of Fen1-mKikGR in our single-molecule assay inhibited cohesin transfer. High concentrations of Fen1-mKikGR result in PCNA being retained on DNA during replication^24^. Because PCNA has been implicated in cohesion establishment^29^, we omitted Fen1-mKikGR in our assays to exclude the possibility of Fen1-mKikGR preventing cohesin transfer due to improper PCNA retention. We visualised the replisome directly using a method similar to one previously described^30^. Endogenous GINS was immunodepleted from *Xenopus* egg extracts and purified fluorescent GINS was used to rescue replication of DNA in GINS-depleted extracts (Supplementary Fig. 5). During replication of λ DNA from single origins, fluorescent CMG moved at the tip of Fen1-mKikGR tracts^30–32^ at an average speed consistent with previous work (426 bp/minute, Supplementary Fig. 6a-c). We next visualised the outcomes of collisions between fluorescent replisomes and pre-loaded JF646-cohesin (Fig. 1a-d, Supplementary Fig. 6d-f). Under these conditions, cohesin transfer was still very rare (5% of events, Fig. 1e) and cohesin sliding ahead of the replisome predominated (67% of events). Therefore, the low frequency of cohesin transfer observed in our previous experiments was not caused by Fen1-mKikGR.

**Figure 1.**
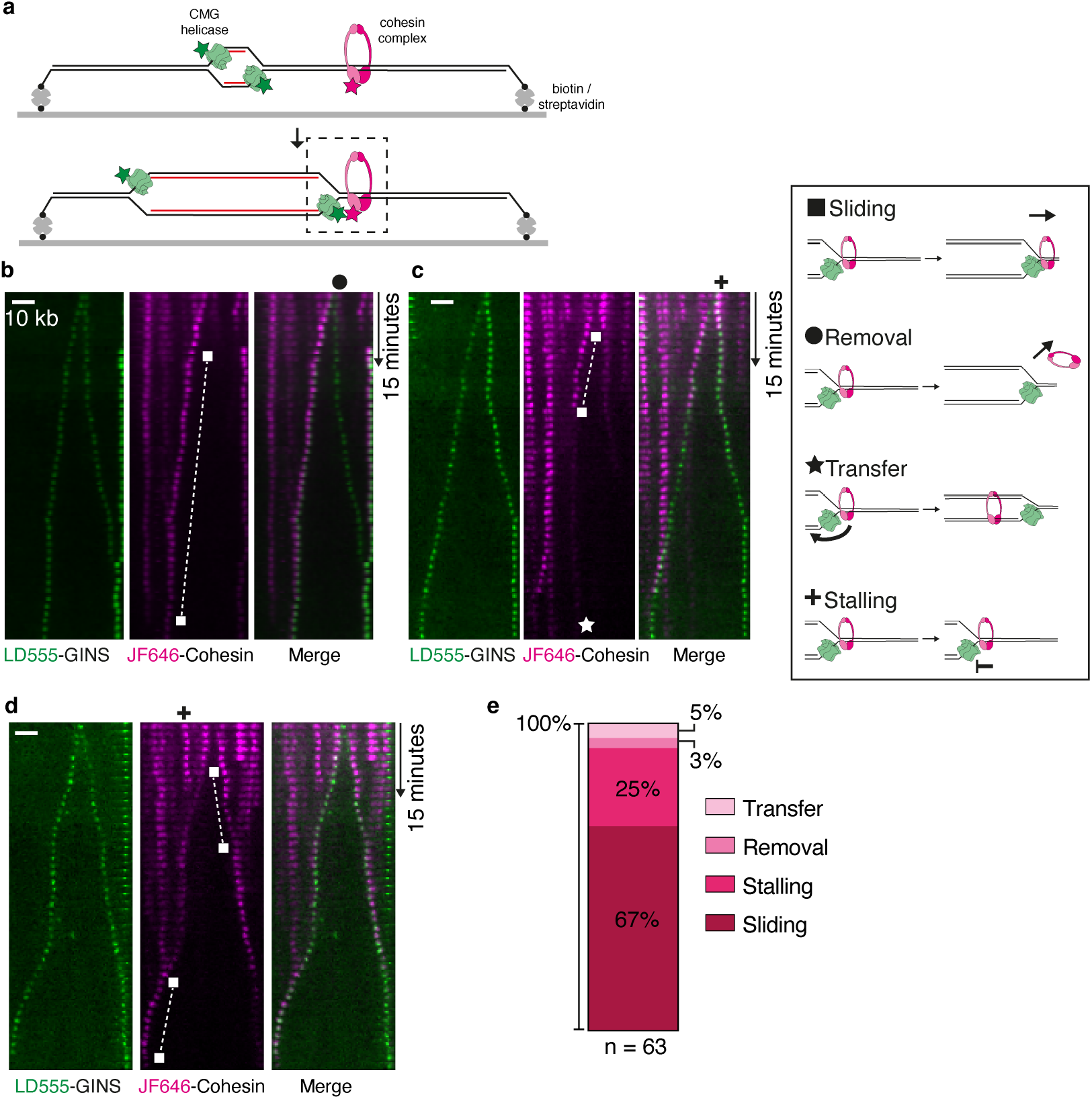
Labelled replisomes push cohesin during DNA replication. **a**, Cartoon showing DNA replication from a single origin with a replisome containing labelled GINS protein. Labelled cohesin is pre-loaded on DNA, and collisions between replisomes and cohesin (dashed box) are visualised. **b**-**d**, Representative kymograms showing LD555-GINS collision with JF646-cohesin. Examples of different cohesin fates (sliding, removal, transfer and fork stalling) are marked with symbols. **e**, Proportions of cohesin fates after collision by labelled GINS complexes.

### Cohesin remains at sites of DNA replication termination

If pre-loaded cohesin is not transferred behind the replisome onto the replicated sister DNAs, how does conversion generate cohesion? We speculated that cohesin pushed ahead of the replisome could generate cohesion when meeting a converging replisome, an event not observed in our previous experiments because they were performed under conditions where origin firing was limited to one per DNA. To determine the fate of cohesin during fork convergence, replication was instead started from multiple origins (Supplementary Fig. 7a). Converging replication forks were visualised using Fen1-mKikGR (Fig. 2a-b, Supplementary Fig. 7b-f). Intriguingly, cohesin pushed ahead of replication forks remained in positions where converging replication forks met. Thus, in 57% of cases, cohesin remained associated with DNA during fork convergence (Fig. 2c), while in 28% of cases, fork convergence was accompanied by cohesin eviction, and in 4% of cases cohesin continued moving after fork convergenc. We observed that cohesin remaining on DNA after fork convergence was resistant to a high-salt wash (HSW) that removed Fen1 from DNAs (Fig. 2a,b, Supplementary Fig. 7b,c), suggesting that this population of cohesin was topologically bound to at least one of the newly replicated DNA molecules^33^.

**Figure 2.**
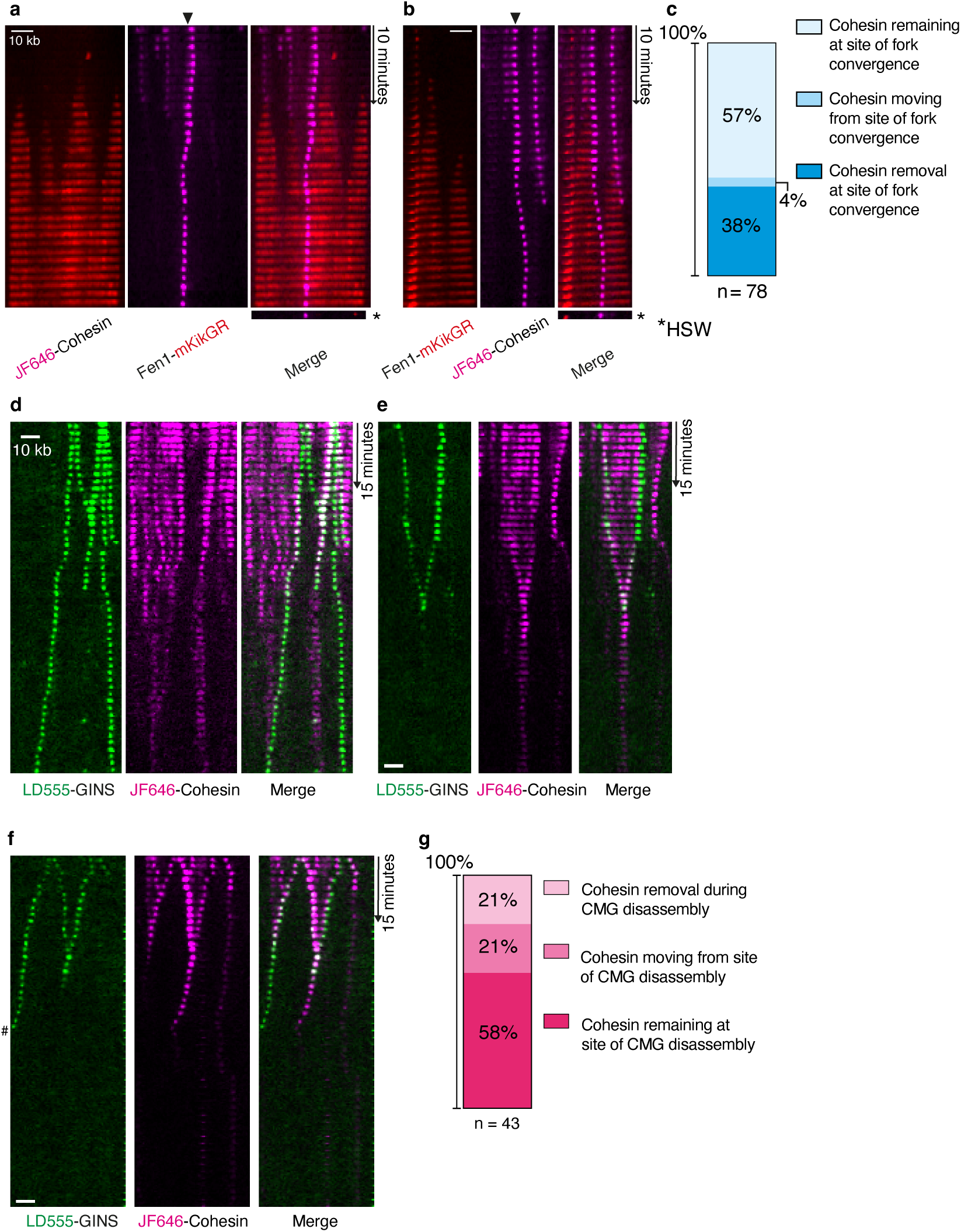
Cohesin is pushed to positions of DNA replication termination. **a, b**, Kymograms showing Fen1-mKikGR labelled replication forks colliding with JF646-cohesin complexes under conditions of high origin firing. After a period of replication, a high salt wash (HSW) was performed and the same λ DNAs were imaged. While Fen1-mKikGR dissociates from DNA, cohesin remains after HSW in these examples. **c**, Quantification of cohesin fates at converging replication forks. The fate of cohesin that was pushed to a converging replication fork or was positioned prior to replication where replication forks converge was measured. **d**-**f**, Kymogram examples showing replisome (LD555-GINS) progression on DNA from multiple origins and colliding with pre-loaded JF646-cohesin. ^#^Indicates where DNA tether snaps before end of kymogram. **g**, Quantification of JF646-cohesin fate at sites where converging replisomes (LD555-GINS) terminate and replisomes are removed.

We envisaged that upon fork convergence, replisomes disassemble while cohesin traps both daughter strands together, thus establishing cohesion at termination sites. To test this hypothesis, multiple origin firing experiments were performed with fluorescent CMG. As expected, replisomes were disassembled shortly after fork convergence^29^ (Supplementary Fig. 8a), even when labelled cohesin persisted on DNA (Fig. 2d-f, Supplementary Fig. 8b-e). Strikingly, in 58% of cases, cohesin remained at the site of replisome disassembly (Fig. 2g). We conclude that cohesin complexes are pushed by advancing replisomes to sites of fork convergence remain at these sites even after replisome disassembly. These findings raise the possibility that pre-loaded cohesin complexes establish cohesion at DNA replication termination sites. The key question is therefore: do the cohesin rings that persist on DNA after replication termination in our assay mediate cohesion?

### Cohesin tethers DNA strands together after DNA replication completion

To assess if cohesin molecules retained at replication termination sites provide cohesion, we developed an assay to measure sister DNA cohesion. Experiments using long DNAs tethered to streptavidin-functionalised surfaces only via 3’-biotins have shown that when a replisome reaches the 5’ DNA end, the new sister DNA that does not contain 3’-biotin is liberated from the surface^30^. The liberated sister DNA collapses and travels with a replisome moving to the opposite end of the template (Supplementary Fig. 9a)^30^. Using this knowledge, we designed a DNA template to visualise interaction between DNA strands after replication (Fig. 3a). We engineered a linear DNA template that contains 48x*lacO* repeats at each end. Binding of Alexa Fluor 488 labelled LacI (LacI-AF488, Supplementary Fig. 9b) to the *lacO* repeats (Supplementary Fig. 9c) prevents replisomes from reaching the DNA ends and the ensuing replicated sister DNA collapse (Fig. 3a). Subsequent removal of LacI-AF488 by IPTG addition results in synchronous completion of replication and collapse of both replicated DNA molecules (Fig. 3a). Importantly, this set up enables us to measure cohesion between the replicated sister DNAs. Lack of cohesion between the replicated DNAs would result in immediate separation of the two sister DNA molecules and their collapse to the respective tethered ends (Fig. 3a, top scenario). If, however the replicated DNAs were held together, the two collapsed sister DNAs would colocalise, as illustrated in Fig. 3a (bottom scenario).

**Figure 3.**
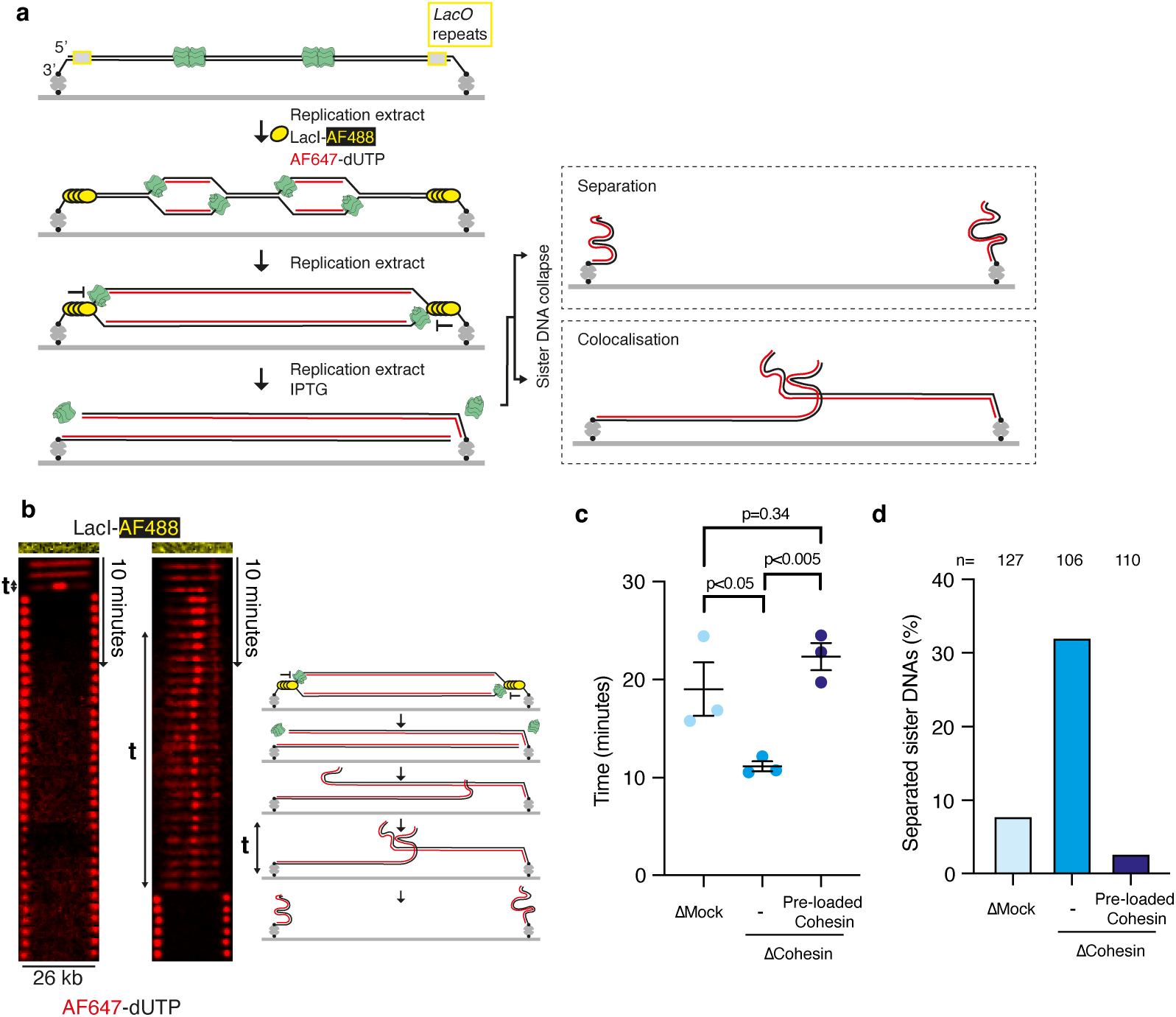
Surface-tethered DNAs have newly replicated sister DNAs bound together by cohesin. **a**, Diagram showing set-up for strand collapse experiments. A 26-kb DNA is tethered exclusively at 3′-ends via biotin-streptavidin interactions to a functionalised glass coverslip. After licensing DNAs with pre-RCs, a replication extract containing LacI-AF488 and AF647-dUTP is added. Most replication forks are paused at DNA ends by bound LacI-AF488. For imaging, excess LacI-AF488 and AF647-dUTP is washed away, then a replication extract containing IPTG is used to remove LacI-AF488 from DNAs. Replication forks then complete replication and sister DNAs collapse is visualised. **b**, Example kymograms where both sister DNA strands collapse as depicted in a cartoon on the right-hand side. The time that collapsed DNA strands remain together is indicated. **c**, Time that collapsed DNA strands remain together in extracts that are mock-depleted, cohesin-depleted and cohesin-depleted with purified cohesin pre-loaded onto DNAs. n=3 independent experiments. Data are mean ± SEM, compared with a two-sided *t-*test. **d**, Percentage of sister DNAs that have separated from one another within 2 minutes of collapsing from both ends under different extract depletion conditions.

We initiated replication from multiple origins in the presence of LacI-AF488, and Alexa Fluor 647-dUTP (AF647-dUTP) was incorporated into nascent DNA for visualisation. Excess LacI-AF488 and AF647-dUTP were washed away and replication extract containing IPTG was added to release LacI-AF488 from DNA ends. Under these conditions, regions of AF647-labelled replicated DNA could be visualised during synchronous collapse of new strands from DNA ends. The assay is intrinsically validated by molecules with partially replicated DNA, where the AF647-dUTP-labelled nascent DNA approached only one DNA end at the LacI barrier. In these instances, after removal of LacI, the collapsed sister DNA moved with the replisome towards the opposite end of the DNA (Supplementary Fig. 9d), as observed previously^30^. Crucially, on molecules where the DNA template was fully replicated up to the LacI barrier at both ends, LacI removal caused collapse of sister DNAs from both ends (Supplementary Fig. 9e). When both new sister DNAs collapsed, the collapsing sister DNAs colocalised together for varying length of time before separating, giving a measure of cohesion. The critical question is whether colocalisation of sister DNAs was because of cohesin mediated cohesion.

To test whether cohesin complexes physically tether collapsing sister DNAs in the experiments described above, the assay was performed in either mock-depleted or cohesin-depleted extracts (Supplementary Fig. 10a) and we compared the time that collapsed sister DNAs colocalised (Fig. 3b-d, Supplementary Fig. 10b,c). The collapsed sister DNAs remained associated significantly and reproducibly longer in mock-depleted extracts compared to cohesin-depleted extracts, indicating that cohesin contributes to physical association of daughter strands (Fig. 3c). Importantly, when purified cohesin was pre-loaded onto tethered DNAs before replication in cohesin-depleted extracts, collapsed sister DNAs remained together for periods of time comparable to that in mock-depleted extracts. In other words, pre-loaded cohesin rescued the defect we observed in cohesin-depleted extracts. After sister DNAs collapse, 32% of sister DNAs were separated within 2 minutes in cohesin-depleted extracts versus 3% when cohesin was pre-loaded before replication in depleted extracts (Fig. 3d). These results show that cohesin complexes pre-loaded onto parental DNA physically tether sister DNAs after replication. Taken together with our previous observation that pre-loaded cohesin is predominantly pushed to sites of replication termination and remains at these sites even after replisome disassembly, our data provide strong evidence that cohesion establishment by cohesin conversion occurs during replication termination.

Our model predicts that cohesive cohesin complexes should be dragged by the collapsing sister DNAs and remain associated with both DNA molecules. To test this, sister DNA collapse experiments were performed with JF549-cohesin pre-loaded onto parental DNA and imaged simultaneously with AF647-dUTP. Fluorescent cohesin molecules colocalising exclusively with the collapsed sister DNA molecules (Fig. 4a, top scenario) would further support cohesin-dependent DNA tethering. Otherwise, if labelled cohesin does not tether sister DNAs together but holds onto individual sister DNAs, we would expect cohesin molecules to remain associated also with stretched DNA after the collapse of replicated DNA molecules (Fig. 4a, bottom scenario). Cohesin was observed to move when both sister DNAs collapsed (Fig. 4c, Supplementary Fig. 11a) and when a single sister DNA collapsed (Fig. 4d, Supplementary Fig. 11b). Strikingly, 85% of cohesin molecules moved and colocalised with collapsed sister DNAs (Fig. 4b), while only a small fraction of cohesin remained on stretched DNAs (15%, Supplementary Fig. 11c). We estimate that approximately 70% of cohesins retained on DNA after replication termination bound both DNA molecules (see Methods). The colocalisation of cohesin with positions where the collapsed sister DNAs remain associated reaffirms the notion that cohesion establishment by pre-loaded cohesin complexes happens during replication termination.

**Figure 4.**
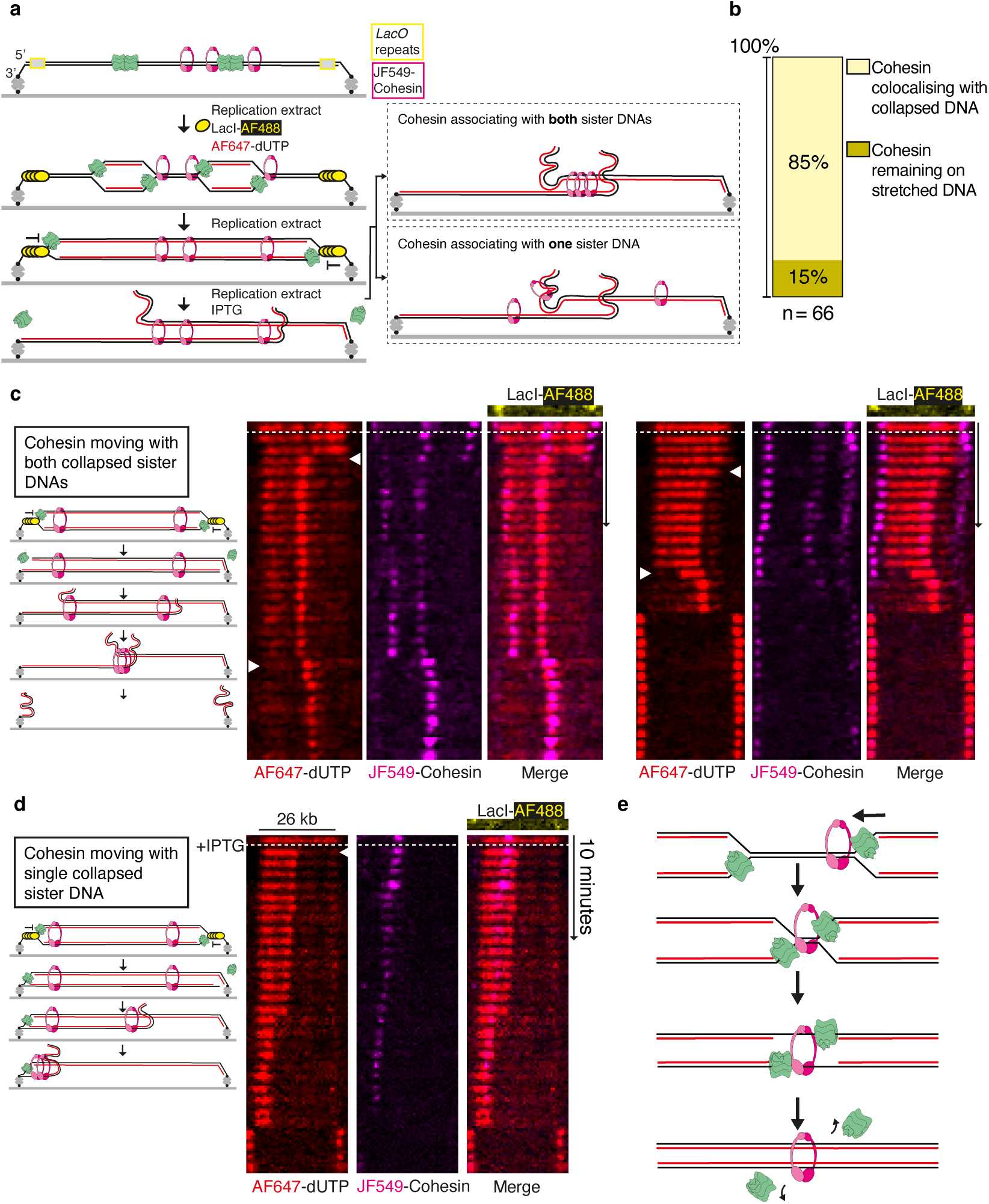
Cohesin binds both sister DNAs of newly replicated surface-tethered DNAs. **a**, Diagram showing the assay used to investigate whether cohesin binds one or both new DNA strands after replication of tethered DNAs in extracts. **b**, Quantification of cohesin position after DNA strand collapse. The position of cohesin, which initially bound AF647-dUTP labelled replicated DNA, upon strand collapse (from either one strand or both strands) was compared. **c**, Kymogram examples where both new DNA strands collapse, and cohesin on replicated DNA remains with the collapsing strands. **d**, Kymogram examples where a single new DNA strand collapses, and cohesin on replicated DNA remains with the collapsing strand. **e**, Model of cohesion establishment during replication termination. Two converging replication forks are depicted in the cartoon. Cohesin is pushed ahead of the leftward moving replisome from the right hand origin. Upon meeting, converging replisomes pull DNA through the topologically loaded cohesin ring to complete DNA replication, so the cohesin ring encircles both new strands of DNA. Replisomes are disassembled.

## Discussion

Prevailing models of cohesion establishment using pre-loaded cohesin propose either passage of the replisome through the cohesin ring or active transfer of cohesin behind the replication fork. According to both scenarios, cohesin is redeposited from parental DNA to newly synthesised sister DNA behind replication forks. We tested this prediction using a single-molecule assay to follow the consequences of encounters between replisomes and pre-loaded cohesin rings. Unexpectedly, we found that cohesin rings are only rarely transferred behind the replisome. Instead, the most frequent outcome of a replisome-cohesin encounter is that pre-loaded cohesin is pushed along the DNA by the advancing replisome. Furthermore, cohesin rings pushed ahead of replisomes to positions of fork convergence are retained on replicated DNA, even after replisome disassembly. Importantly, using newly developed sister DNA collapse assays, we show that cohesin rings that remain at the sites of replication termination are capable of tethering sister DNAs. We note that in our sister DNA collapse assays, when cohesin is holding on collapsed sister DNAs, most collapsed sister DNAs do eventually come apart from one another. This could be explained by the fact that the cohesin rings holding the two linear, relatively short, sister DNAs eventually slide out of the free DNA ends. Indeed, circular minichromosomes that are held together by cohesin in yeast cells lose their cohesion upon linearization of the minichromosome^34^. Loss of cohesin association with circular DNAs *in vitro* upon linearisation is frequently used as a surrogate measure of topological association^35^. The strand collapse assay presented here provides a new way to assess sister chromatid cohesion establishment.

The notion of replisomes pushing cohesin is supported by transcription repositioning cohesin on yeast chromosomes^21^ and cohesin being pushed ahead of T7 RNAP and FtsK *in vitro*^36,37^. A previous work visualising replication fork collision with cohesin in *Xenopus* extracts found that cohesin being pushed by the replisome occurred 15% of the time whilst cohesin was incorporated into replicated DNA in 32% of cases^38^, which on face value would appear inconsistent with our observations. Because this study used firing from multiple origins to replicate the DNAs, we suggest that the cohesin transfer events they observed were largely the result of cohesin being incorporated into replicated DNA during termination. In the limited number of examples where single replication forks encountered cohesin in their experiment, cohesin sliding or eviction events were observed^38^.

Our data strongly support a mechanism where cohesion establishment using pre-loaded cohesin rings happens during replication termination. We provide the first experimental evidence of this idea previously raised as a theoretical model^39^. We propose a simple mechanism whereby cohesion establishment during termination negates the need either for replisomes to pass through cohesin rings or for rings to transiently open during transfer to replicated DNA (Fig. 4e). According to this model, cohesin complexes that have entrapped DNAs are pushed by the replisome to termination sites, whereupon replisomes remain either side the cohesin ring and the final stretches of unreplicated parental DNA could be pulled by the CMG helicase through cohesin rings to complete replication. As replisomes move onto the new double-stranded DNA, they lose interaction with the excluded DNA strand leading to CMG ubiquitylation and replisome disassembly^40^. In this scenario, cohesin rings would end up encircling the two new DNA strands and provide cohesion. Alternative models can be envisaged where structural rearrangements of cohesin and/or the replisome occur specifically at replication termination sites.

TIMELESS/TIPIN, AND1 and DDX11 are the replisome associated proteins involved in converting pre-loaded cohesin to generate cohesion, but how these factors mediate cohesin conversion is unclear^4,19^. As components of the replisome, TIMELESS/TIPIN and AND1 have a general role in maintaining normal replisome speeds during replication of chromatin^41^. TIMELESS/TIPIN and AND1 also recruit the ubiquitin ligase CRL2^LRR1^ during replisome disassembly^40,42^. AND1 binding to *Xenopus* replisomes is highest during replication initiation and termination^43^. DDX11 is a superfamily 2 DNA helicase that interacts specifically with numerous replication factors^44–46^. Single-molecule assays can be used to unravel whether TIMELESS/TIPIN, AND1 and DDX11 are involved in the pathway of cohesin conversion during replication termination we have proposed here. An intriguing possibility is that the roles of these factors in replisome disassembly are directly linked to cohesion establishment.

## Materials and Methods

### Protein expression, purification and labelling

#### Fen1-mKikGR

Purification was carried out as previously described^24^.

#### Cohesin-trimer and tetramer (Halo-JF646/JF549)

Construction of the expression vectors for cohesin tetramer was described previously^15^. To generate vectors containing cohesin trimer, XSMC1 and XSMC3 with a C-terminal Halo and 2X Flag tags were cloned into MultiBac vectors (pACEbac1). XRAD21 with a C-terminal His8 tag was cloned into pDIC plasmid. The pACEbac1 XSMC1 XSMC3-Halo-Flag and pIDC XRAD21-8xHis were then combined by a Cre recombinase reaction (New England Biolabs). To generate the baculoviruses, DNAs were first transformed into DH10Bac (Thermo Fisher) cells and bacmids containing the expression vector screened for by blue-white selection. Bacmid DNA was then extracted and 2 µg of bacmid DNA was transfected into 2 ml *S. frugiperda* Sf9 cells (Thermo Fisher) at a cell density of 1 × 10^6^ cells ml^−1^ using FuGENE HD reagent (Promega), grown in Sf900 II SFM media (Thermo Fisher). These were then incubated at 27°C for 5 days to create P1 virus. P2 virus was then amplified by infecting 50 ml Sf9 cells at a density of 2 × 10^6^ cells ml^−1^ with 500 µl P1 virus and incubating in the dark at 27°C for 3 days with shaking at 100 rpm.

Typically, proteins were expressed by adding 5 ml P2 virus to 500 ml Sf9 cells at a density of 2 × 10^6^ cells ml^−1^ and incubating in the dark at 27°C for 2 days with shaking at 100 rpm. Cells were then harvested by centrifugation at 1000*g*, washed with PBS, and then frozen in liquid nitrogen and stored at −80°C. All subsequent steps were performed on ice or at 4°C. Cells were lysed by thawing and dounce homogenizing in buffer A500 (25 mM HEPES-KOH pH 7.5, 500 mM KCl, 5% v/v glycerol), 20 mM β-mercaptoethanol, 0.05% v/v Tween-20, 0.5 mg/ mL PMSF and complete protease inhibitor (Roche). After lysis, an equal volume of buffer A0 (buffer A500 lacking KCl) was added to the lysate and centrifuged at 75,000*g* for 40 min. The clarified lysate was filtered through a 0.45 µm filter and cohesin purified in the AKTA system using a 5 mL (HisTrap) TALON column (VWR). The column was washed with 20 column volumes of buffer B (20 mM Tris PH7.5, 250 mM KCL, 5% glycerol) and eluted over a linear gradient of 0-500 mM imidazole. The peak fractions containing cohesin were pooled and incubated with anti-FLAG-M2 resin (Sigma) and 10nM JF646 or JF549 HaloTag ligand (a kind gift from Luke Lavis^47^) for 3 hours at 4°C in the dark. The beads were pelleted and washed with 5X EB buffer (100 mM KCl, 2.5 mM MgCl_2_ and 50 mM HEPES-KOH pH 7.5) and eluted in EB buffer with FLAG peptide containing 5% glycerol and 5 mM DTT. Finally, proteins were purified via size exclusion chromatography using a Superose 6 increase 10/300 GL column (VWR). Peak fractions were collected, concentrated to typically 3 µM and stored in aliquots at −80°C.

#### SA1

XSA1 was cloned into pACEbac1 vector with a C-Terminal His8 tag. The P1 and P2 viruses were generated as described above. SA1 was expressed by adding 5 ml P2 virus to 500 ml Sf9 cells at a density of 2 × 10^6^ cells ml^−1^ and incubating in the dark at 27°C for 2 days with shaking at 100 rpm. Cells were then harvested by centrifugation at 1000*g*, washed with PBS, and then frozen in liquid nitrogen and stored at −80°C. All subsequent steps were performed on ice or at 4°C. Cells were lysed by thawing and dounce homogenizing in buffer A500 (25 mM HEPES-KOH pH 7.5, 500 mM KCl, 5% v/v glycerol), 20 mM β-mercaptoethanol, 0.05% v/v Tween-20, 0.5 mg/ mL PMSF and complete protease inhibitor (Roche). After lysis, an equal volume of buffer A0 (buffer A500 lacking KCl) was added to the lysate and centrifuged at 75,000*g* for 40 min. The clarified lysate was filtered through a 0.45 µm filter and cohesin purified in the AKTA system using a 5 mL (HisTrap) TALON column (VWR). The column was washed with 20 column volumes of buffer B (20 mM Tris PH7.5, 250 mM KCL, 5% glycerol) and eluted over a linear gradient of 0-500 mM imidazole. The peak fractions containing SA1 were pooled concentrated to 1ml final volume and was purified via size exclusion chromatography using a Superose 6 increase 10/300 GL column (VWR). Peak fractions were collected, concentrated to typically 3 µM and stored in aliquots at −80°C.

#### NIPBL-C

HsNIPBL-C (residues 1163-2804) was cloned into pAECbac1 vector with an N-terminal His6 and a C-terminal Flag tag. P1 and P2 viruses were generated as described above. NIPBL-C was expressed by adding 10 ml P2 virus to 500 ml Sf9 cells at a density of 2 × 10^6^ cells ml^−1^ and incubating in the dark at 27°C for 2 days with shaking at 100 rpm. Cells were then harvested by centrifugation at 1000*g*, washed with PBS, and then frozen in liquid nitrogen and stored at −80°C. All subsequent steps were performed on ice or at 4°C. Cells were lysed by thawing and dounce homogenizing in buffer A500 (25 mM HEPES-KOH pH 7.5, 500 mM KCl, 5% v/v glycerol), 20 mM β-mercaptoethanol, 0.05% v/v Tween-20, 0.5 mg/ mL PMSF and complete protease inhibitor (Roche). After lysis, an equal volume of buffer A0 (buffer A500 lacking KCl) was added to the lysate and centrifuged at 75,000*g* for 40 min. The clarified lysate was filtered through a 0.45 µm filter and cohesin purified in the AKTA system using a 5 mL (HisTrap) TALON column (VWR). The column was washed with 20 column volumes of buffer B (20 mM Tris PH7.5, 250 mM KCL, 5% glycerol) and eluted over a linear gradient of 0-500 mM imidazole. The peak fractions containing NIPBL were pooled and incubated with anti-FLAG-M2 resin (Sigma) and for 3 h at 4°C. The beads were pelleted and washed with 5X EB buffer (100 mM KCl, 2.5 mM MgCl_2_ and 50 mM HEPES-KOH pH 7.5) and eluted in EB buffer with Flag peptide containing 5% glycerol and 5 mM DTT. The protein was concentrated to typically 1 µM and stored in aliquots at −80°C.

#### GINS

Purification was adapted from a previously described protocol^30^. The MultiBac vectors pFL-Psf1/Psf2 and pSPL-Sld5/Psf3-LPETG-Flag were cloned using codon-optimised GINS sequences synthesised by GeneArt (Sigma). Sequences encoding a linker (GGGGSGGGGS), a Sortase labelling tag (LPETG) and a Flag epitope (DYKDDDDK) were included to be incorporated at the C-terminus of Psf3. pFL-Psf1/Psf2 and pSPL-Sld5/Psf3 plasmids were combined by Cre-Lox recombination, and the fused pFL/pSPL plasmid transformed into DH10MultiBac cells to create a bacmid. Sf9 insect cells were transfected and multiple rounds of virus amplification were performed. Hi5 insect cells (
1 × 10^6^ cells/mL) were infected for GINS expression and were harvested after 48 hours.

Purification steps were performed at 4°C. A pellet from 1 L cells was resuspended in 40 mL lysis buffer (20 mM Tris pH 8, 500 mM NaCl, 0.1% NP40, 10% glycerol, 2 mM β-mercaptethanol, 1 x EDTA-free protease inhibitor tablet (Roche)) and sonicated for 2 min. The lysate was cleared with centrifugation at 30,000*g* for 30 min, and the supernatant incubated with 3 mL pre-washed Anti-FLAG M2 affinity gel (Sigma) for 2 hours. The resin was washed 2 × 15 mL wash buffer (20 mM Tris pH 8, 500 mM NaCl, 0.1% NP40, 10% glycerol and 2 mM β-mercaptethanol). Protein was eluted with 2 × 5 mL wash buffer/333 µg/mL Flag peptide. The Flag resin eluate was loaded onto a MonoQ 5/50 GL (Cytiva) column prequilibriated with MonoQ-100 (20 mM Tris pH 7.5, 100 mM NaCl, 10% glycerol and 1 mM DTT). GINS was eluted using a linear gradient of MonoQ-700 (20 mM Tris pH 7.5, 700 mM NaCl, 10% glycerol and 1 mM DTT).

For fluorescent labelling, 3 volumes of a mixture containing 4 nmol GINS, 1 nmol Sortase A and 100 nmol fluorescent peptide (NH2-GGGHHHHHHC(*)-COOH, where *=LD555, LD655 or AF647, conjugated using maleimide-thiol reaction) was added to 1 volume of labelling buffer (80 mM HEPES pH 7.5, 600 mM NaCl, 40% glycerol, 20 mM CaCl_2_ and 4 mM DTT). After an overnight reaction, GINS was purified using a Sepharose 200 10/300 GL size exclusion column equilibrated in GF buffer (20 mM Tris pH 7.5, 150 mM NaCl, 10% glycerol and 1 mM DTT). To selectively purify labelled GINS containing polyhistidine, peak fractions were supplemented with 10 mM imidazole and 10 µg/mL aprotinin/leupeptin and bound to NiNTA beads (Qiagen) for 2 hours. Beads were washed with GF buffer/10 mM imidazole then GINS eluted with GF buffer/500 mM imidazole. The protein was dialysed in GF buffer overnight before storage at −80°C.

#### LacI-AF488

Labelled LacI was purified essentially as described^48^. Plasmids containing LacI with a C-terminal HHHHHHC (‘LacI-Far’, a gift from Sebastian Deindl^48^) was transformed into BL21 cells. Expression was induced for 3 hours in 1 L of cells using 0.2% L-rhamnose. Cell pellets were resuspended in 20 mL buffer A (20 mM phosphate pH 7.4, 500 mM NaCl, 5 mM β-mercaptoethanol, 20 mM imidazole) with 1 EDTA-free protease inhibitor tablet (Roche) and supplemented with 10 µg/mL lysozyme and 25 U/mL benzonase. After sonication for 2 min total, lysate was cleared by centrifugation at 20,000*g* for 20 min and passed through a 0.45 µm filter. LacI was bound to a 5 mL HisTrap HP column equilibrated in buffer A and eluted with a linear gradient of buffer B (20 mM phosphate pH 7.4, 500 mM NaCl, 5 mM β-mercaptoethanol, 500 mM imidazole). Protein was buffer exchanged into phosphate buffered saline (PBS) containing 10% glycerol using a spin concentrator (10 kDa MWCO, Millipore).

For labelling, 1 mg Alexa Flour 488 C_5_ Maleimide (ThermoFisher) was dissolved in 50 µL DMSO then mixed with ~0.5 µmol LacI in degassed PBS containing 10% glycerol and 500 µM TCEP. After 90 min at room temperature, the reaction was quenched with 5 mM β-mercaptoethanol. LacI-AF488 was purified with a 5 mL HisTrap HP column (as described above), before peak fractions were dialysed into PBS with 500 mM NaCl overnight. LacI-AF488 was diluted 1:1 with glycerol before storing aliquots at −80°C.

### Xenopus laevis egg extract preparation and immunodepletion

#### Xenopus laevis egg extract preparation

Animal husbandry, injections and egg collection were performed by the Francis Crick Institute Aquatics Facility. Extracts and sperm chromatin were prepared as described^49^ and aliquots stored at −80°C.

#### Cohesin immunodepletion

For depletions, rProtein A Sepharose Fast Flow (PAS, Cytiva) beads were extensively washed with PBS before antibody binding. After antibody binding, beads were washed 3 times with PBS and 5 times with Egg Lysis Buffer (ELB, 50 mM HEPES pH 7.7, 2.5 mM MgCl_2_, 50 mM KCl), before transferring to siliconised microcentrifuge tubes. To recover extracts from PAS beads between rounds of depletion, the extract/bead mixture was applied to a homemade nitex filter^49^ and centrifuged at 2,800*g* for 40 seconds.

SMC1 and SMC3 antibodies were a kind gift from Vincenzo Costanzo. Both antibodies were raised in rabbits immunised with peptides: anti-SMC1 with DLTKYPDANPNPND and anti-SMC3 with EQAKDFVEDDTTHG. A cysteine was added to the N-terminus of both peptides, and the modified peptides were used for immunoaffinity purification according to manufacturer’s instructions (SulfoLink Immobilization Kit for Peptides, ThermoFisher). 72 µL of PAS beads was incubated overnight with 100 µg purified SMC1 antibody and 100 µg purified SMC3 antibody. 1 µL 0.5 mg/mL nocodazole was added to 60 µL HSS and 60 µL NPE supplemented with DTT to a final concentration of 10 mM. HSS and NPE were mixed and added to 24 µL of anti-SMC1/anti-SMC3 PAS beads, for 3 rounds of depletion for 45 min at 4°C. For cohesin depletions used for strand collapse experiments, the same ratios of SMC1/SMC3 antibodies to beads were used. Mock depletions used PAS beads washed with ELB only. 60 µL HSS was added to 10 µL beads. A mixture of 90 µL HSS with 90 µL NPE was added to 30 µL PAS beads. Depleted extract was aliquoted and frozen at-80°C before use.

#### GINS immunodepletion

Rabbits were immunised with *Xl*GINS purified from insect cells. Anti-GINS antibody was affinity purified using protein A-sepharose (Covalab). For bulk replication assays, HSS and NPE were separately depleted. 300 µL purified anti-GINS antibody (3.5 mg/mL) was incubated with 120 µL PAS beads overnight. 180 µL HSS was supplemented with 3 µL 0.5 mg/mL nocodazole and added to 30 µL anti-GINS PAS beads for 2 rounds of depletion for 45 min at 4°C. 90 µL of NPE supplemented with DTT to a final concentration of 10 mM was added to 20 µL anti-GINS PAS beads for 3 rounds of depletion. 15 µL ΔGINS extracts were stored at −80°C before use.

For single-molecule replication assays, a mixture of HSS and NPE was depleted. 150 µL purified anti-GINS antibody (3.5 mg/mL) was incubated with 40.5 µL PAS beads overnight. 13.5 µL of washed beads were used for each round of depletion. 22.5 µL of HSS was supplemented with 0.35 µL 0.5 mg/mL nocodazole and mixed with 45 µL of NPE supplemented with DTT to a final concentration of 10 mM. The extract mixture was depleted for 3 rounds of 1 hour, before storing 15 µL aliquots of HSS/NPEΔGINS at −80°C.

### Preparation of DNA substrates

#### mBiotin-λ-mDigoxigenin

PCR reactions were used to incorporate biotin- or digoxigenin-modified nucleotides into handles ligated to the ends of λ DNA. Two PCR reactions using GoTaq G2 PCR mix (Promega) and a pUC19 template were set up: PCR-Dig with oGC101/oGC102 primers and 25 µM digoxigenin-11-dUTP (Roche), and PCR-Bio with oGC101/oGC103 primers and 25 µM biotin-16-dUTP (Enzo). The products were isolated with a PCR purification kit (Qiagen), nicked with Nt.BspQI (NEB) and heated to 65°C to create 12 bp ssDNA ends complementary to λ DNA ends. Before cooling, PCR-Bio was mixed with oGC104 and PCR-Dig was mixed with oGC105 to prevent reannealing. The handles were separated on a 1.5% agarose gel and purified using a gel purification kit (Qiagen). λ DNA was phosphorylated with T4 polynucleotide kinase (NEB), ligated to PCR-Bio/Nt.BspQI with T4 DNA ligase (NEB) and purified from a 0.5% agarose gel by electroelution. The product was ligated to PCR-Dig/Nt.BspQI and purified once more, before aliquots were stored at − 20°C and freeze-thaw cycles avoided.

#### List of oligonucleotides used for making mBiotin-λ-mDigoxigenin substrates

*oGC101 ATGCCGGGAGCAGACAAGCCCGTC*

*oGC102*

*GGGCGGCGACCTGGAAGAGCAGCTGGCACGACAGGTTTCCCG*

*oGC103*

*AGGTCGCCGCCCGGAAGAGCAGCTGGCACGACAGGTTTCCCG*

*oGC104 AGGTCGCCGCCC*

*oGC105 GGGCGGCGACCT*

#### Biotin-λ-Biotin

Doubly biotinylated DNA was prepared as previously described^28^. 10 µg λ DNA (NEB) was added to a 50 µL reaction with 80 µM biotin-dCTP (Invitrogen), 80 µM biotin-dATP (Invitrogen), 100 µM dTTP and 100 µM dGTP, then heated to 65°C for 5 min. 2.5 U Klenow polymerase (NEB) was added and the reaction was incubated for 30 min at 37°C for 30 min, then at 70°C for 15 min. DNA was purified by electroelution from a 0.5% TBE gel, then dialysed into 10 mM Tris pH 7.5 and 1 mM EDTA. Aliquots were stored at −20°C and freeze-thaw cycles avoided.

#### Biotin-LacO-23kb-LacO-Biotin

pHY42, a 17.3 kb plasmid containing a 48x*lacO* array (~1.5 kb), was digested with BsiWI-HF restriction enzyme. The 4 bp overhangs formed were filled in with biotin-11-dGTP (Jena Bioscience), biotin-16-dUTP (Roche), biotin-14-dCTP (Invitrogen) and biotin-14-dATP (Invitrogen), each added to a final concentration of 50 µM in a reaction with ~6 µg DNA and 15 U Klenow polymerase. The reaction was buffer exchanged in a Microspin G-50 Column (Cytiva) before treatment with Quick Calf Intestinal Phosphatase (CIP, NEB). The plasmid was digested with AgeI, creating a 12.9 kb fragment that was separated from a smaller 4.4 kb fragment on a 0.6% agarose gel, then purified by electroelution. The fragment was self-ligated with T4 DNA ligase (NEB) to create a 25.8kb linear DNA with a 48x*lacO* array and 3’-biotins at either end. The DNA substrate was purified from a 0.5% agarose gel by electroelution.

### Coverslip functionalisation, microfluidic flow channel preparation and DNA tethering

Coverslips were functionalised and microfluidic flow channels prepared essentially as previously described^25^. 24 × 60 mm glass coverslips (VWR) were sonicated in ethanol for 30 min and 1 M KOH for 30 min, with rinsing in water performed between sonications. This was repeated once before plasma cleaning and silane treatment in 2% v/v 3-aminopropyltriethoxysilane (in acetone) for 2 min. After rinsing in water, 75 mg mPEG-SVA (MW 5,000, Laysan Bio) and 2 mg biotin-PEG (MW 5,000, Laysan Bio) dissolved in 500 µL 100 mM NaHCO_3_ was placed between two coverslips and incubated for 3 hours. Coverslips were rinsed and stored under vacuum.

Microfluidic flow channels were assembled using a cut glass slide with holes drilled for PE20 inlet and PE60 outlet (Intramedic) polyethylene tubing. Double sided tape (AR90880, Adhesive Research) was cut and sandwiched between the coverslip and glass slide, creating a flow channel sealed with epoxy resin.

Flow channel outlet tubing was connected to a syringe pump (Harvard Apparatus) and the flow channel washed in blocking buffer (20 mM Tris pH 7.5, 50 mM NaCl, 2 mM EDTA, 0.2 mg/mL BSA). 0.2 mg/mL streptavidin in blocking buffer was incubated in the flow channel for 10 min before DNA tethering. mBiotin-λ-mDigoxigenin DNAs were diluted to <1 ng/µL in blocking buffer and incubated for up to 30 min to tether the biotinylated end then washed. Single-tethered DNA was stretched with blocking buffer containing 1 µg/mL biotinylated anti-digoxigenin (Perkin Elmer) at 100 µL/min flow rate. Biotin-λ-biotin and Biotin-LacO-23kb-LacO-Biotin DNAs, at a concentration <1 ng/µL, were bound to surfaces at 100 µL/min flow rate for between 2 and 10 min. Tethered DNAs were stained with 5 nM SYTOX Orange (ThermoFisher) to check DNA concentration and DNA end-to-end distances.

### Cohesin loading on chromatin

HSS extracts were supplemented with either EB buffer (100 mM KCl, 2.5 mM MgCl_2_ and 50 mM HEPES-KOH pH 7.5) or with 60nM Geminin and incubated at for 23 °C for 10 min. This was followed by addition of 400nM recombinant cohesin and sperm chromatin. Reactions were incubated at 23 °C for 60 min. To isolate HSS extract assembled chromatin, samples were diluted in ten volumes of EB buffer containing 0.25% Nonidet P-40 and centrifuged through a 30% sucrose (in EB) layer at 10,000 rpm for 5 min at 4°C using a HB-6 rotor (Sorvall), washed three times with 500 µL EB buffer and centrifuged at 10,000 rpm for 1 min. The pellet was resuspended in Laemmli loading buffer and the proteins resolved by either 4%–15%, 7.5% or 10% SDS-PAGE and analyzed by western blotting with specific antibodies as indicated.

### Cohesin loading onto tethered DNAs and DNA replication with labelled Fen1

Prior to DNA replication with Fen1-mKikGR-labelled replication bubbles, labelled cohesin was loaded onto mBiotin-λ-mDigoxigenin or Biotin-λ-Biotin template DNAs during replication licensing. An ATP regeneration mixture was assembled with 5 µL 0.2 M ATP, 10 µL 1 M phosphocreatine and 0.5 µL 25 U/mL creatine phosphokinase. ELBS buffer used to dilute extracts was made from 30 µL ELB containing 0.25 M sucrose and supplemented with 1 µL ATP regeneration mixture. 33 µL HSS was mixed with 0.5 µL 0.5 mg/mL nocodazole and 1 µL ATP regeneration mixture. 16 µL NPE was mixed with 0.5 µL ATP regeneration mixture, and both HSS and NPE extracts were centrifuged for 5 min at 16,000*g* before use. A licensing mixture was assembled containing 15 µL HSS, 15 µL ELBS and 0.75 µL 400 ng/µL oligonucleotide duplex^25^, and JF646-labelled cohesin tetramer was added to a final concentration of 500 nM. DNA was incubated with licensing mixture for 25 min. A mixture of 15 µL NPE, 15 µL HSS and 15 µL ELBS supplemented with 5 ng/µL pBlueScript and 2.5 µM Fen1-mKikGR was split into two, with half the mixture infused into flow channels to initiate DNA replication. The remainder was supplemented with 0.1 µg/mL p27^kip^ and added to flow channels after between 2 and 10 min to limit further origin firing. For experiments with cohesin-depleted extract, 30 µL of cohesin-depleted HSS/NPE was mixed with 1 µL ATP regeneration mixture. The depleted extract was then mixed with 15 µL ELBS and supplemented with pBlueScript and Fen1-mKikGR as described above. For imaging in high salt buffer after replication, ELB containing 500 mM KCl was infused into the flow channel.

### DNA replication with labelled GINS

#### Bulk DNA replication assay with GINS-depleted extracts

HSS and NPE were prepared as described above, with NPE supplemented with [α^32^P]-dATP. 10 ng/µL pBlueScript was added to HSS and licensing performed for 30 min. 1 volume licensing mixture was added to 1 volume 1.5 µM GINS or 0.3 µM AF647-GINS diluted in ELBS, then 1 volume NPE was added to begin DNA replication. Reactions were stopped and separated on a 0.8% agarose gel before visualisation by autoradiography.

#### Single-molecule replication assay with GINS-depleted extracts

Biotin-λ-Biotin template DNAs were used for reactions with labelled GINS. HSS and NPE were prepared as above. 15 µL GINS-depleted HSS/NPE was mixed with 0.5 µL ATP regeneration mixture. 10 µL HSS, 10 µL ELBS and 0.5 µL 400 ng/µL oligonucleotide duplex were mixed and infused into flow channels for 10 min of licensing. The channel was washed with 60 µL ELB supplemented with 1 mg/mL BSA and 0.5 mg/mL casein. A mixture of 15 µL GINS-depleted HSS/NPE, 5 µL ELBS, 4 µL LD655-/LD555-GINS (~2-5 µM), 0.5 µL 150 ng/µL pBlueScript and 0.2-1 µL 0.5 mg/mL cyclin A2 (Abcam) was infused into the flow channel for 2-5 min. To prevent further origin firing and wash away excess fluorescent GINS, a mixture of 15 µL NPE, 22 µL HSS, 15 µL ELBS, 2 µL 0.1 µg/mL p27^kip^ and 1.2 µL 150 ng/µL pBlueScript, supplemented with ~2.5 µM Fen1-mKikGR when required, is added to the flow channel.

To load cohesin on DNA before replication with labelled GINS, cohesin was loaded onto DNAs in buffer prior to licensing. 1 µL 2 µM JF646-cohesin trimer (Smc1/Smc3/Rad21), 1.5 µL 3 µM SA1 and 1.5 µL 3 µM NIPBLc were mixed and incubated on ice for 10 min. Cohesin buffer was made from ELB supplemented with 1 mg/mL BSA, 5 mM DTT, 0.002% Tween-20 and 5% glycerol, and this was used to wash DNAs tethered in flow channels. 1 µL cohesin/loader mixture was diluted in 200 µL cohesin buffer containing 3 mM ATP, and incubated with tethered DNAs for 10-15 min. The flow channel was washed with cohesin buffer before replication with labelled GINS as described above.

### Experiment to monitor DNA strand collapse

For experiments with labelled cohesin pre-loaded onto DNAs, JF549-cohesin was loaded onto tethered *Biotin-LacO-23kb-LacO-Biotin* DNAs in buffer as described above. HSS and NPE were prepared as above. Licensing was performed with a mixture of 15 µL HSS, 5 µL ELBS and 0.75-1.5 µL 400 ng/µL oligonucleotide duplex for 10 min. 30 µL NPE, 30 µL HSS, 30 µL ELBS and 2-4 µL 150 ng/µL pBlueScript were mixed and split into 3 × 30 µL reaction mixtures. The first firing extract was supplemented with 2-3 µL 0.5 mg/mL cyclin A2, 1.5-1.8 µL 1 mM AF647-dUTP (Jena BioScience) and 0.75 µL 25 µM LacI-AF488 and incubated in the flow channel for between 10 and 15 min. The second firing mixture was infused into the flow channel to remove excess AF647-dUTP and LacI-AF488 for 3 min. A third firing mixture was supplemented with 1.5 µL 1 M IPTG, to remove LacI from the *lacO* sequences at the ends of DNA, was infused into the flow channel.

To compare the length of time collapsed DNA strands survived in differently depleted extracts, experiments were essentially performed as described above. Licensing mixture was supplemented with 1.5 µL 400 ng/µL oligonucleotide duplex and 3 µL 0.2 mg/mL Cdt1^50^ to ensure maximal pre-RC assembly. To remove as much fluorescent nucleotide as possible, the volume of the third firing mixture was increased to 60 µL and infused at a 1.5 µL/min flow rate for 40 min. Images were taken for 60 min for each type of depleted extract.

### Image acquisition and analysis

#### Image acquisition and processing

Images were acquired using a Nikon Eclipse Ti microscope as previously described^28^. A 5×5 or 6×6 grid of field of view was imaged during DNA replication, typically with a lapse time of 60-90 seconds. Images were initially processed in NIS Elements, with the “Advanced Denoising” function used with a denoising power of 5 for each channel. In some cases, a rolling ball background correction (radius 0.96 µm) was used for background subtraction. Images were corrected for drift using the align function. Fiji was then used rotate and crop regions of interest, with a width of 5-7 pixels, and to create kymograms using the “Make Montage” function.

#### Measuring fork speeds

Average DNA lengths were measured for each experiment in BB supplemented with 5 nM SYTOX Orange without any flow. To calculate fork speeds, Fen1-mKikGR replication bubble growth was measured during a period of constant fork movement and averaged between 2 diverging replisomes. The rates of individual labelled replisome molecules were measured individually.

#### Defining cohesin fates after collision with the replication forks

Cohesin-fork encounter was defined as colocalisation of cohesin signal (diffraction-limited spot) with the tip of Fen1-mKikGR-decorated replication bubble. Cohesin removal was marked by the loss of cohesin fluorescence in the next time frame upon fork encounter. Cohesin transfer was assigned when, upon fork encounter, cohesin signal was incorporated into the replication bubble and could be followed for at least two subsequent time frames (2 min). Cohesin sliding was determined by a unified cohesin-fork movement over at least 3 pixels. Replication fork stalling was assigned if a fork movement was arrested by a static (within 2 pixels) cohesin fluorescence for at least three time frames (3 min).

#### Defining cohesin fates after collision by labelled replisomes

When the replisome and cohesin co-localise in a diffraction limited spot moving >2 pixels in <5 min, cohesin sliding was scored. When co-localising replisomes and cohesin do not move >2 pixels in >5 min, stalling was scored. When there was no detectable change in replisome speed during co-localisation with cohesin, and cohesin fluorescence is lost without moving >2 pixels, the event was defined as removal. When a replisome and cohesin co-localise for <5 min, and cohesin remains in the same spot whilst the replisome moves >2 pixels away, a transfer event was scored. Cohesin fates during replication termination were defined similarly. Only events where both converging replisomes were labelled, at least one of these converging replisomes is associated with a sliding cohesin and where the replisome is disassembled after convergence were included.

#### Strand collapse experiments

Cohesin and the collapsed strand were considered to be colocalised when <2 pixels apart. We assumed the probability of cohesin binding exclusively to either strand was equal. 15% of cohesin colocalised with the stretched DNA strand so we assumed a further 15% of cohesin colocalised exclusively with the collapsed DNA strand, therefore inferring that 70% of cohesin bound to both DNA strands.

To compare the length of time collapsed DNA strands survived in differently depleted extracts, only events where the entire DNA was replicated and both DNA strands collapsed were considered. Events where collapsed DNA strands remain together until the end of 60 min imaging were also included.

## Acknowledgements

We thank the Francis Crick Institute Aquatics Facility for *Xenopus laevis* husbandry and egg collection. We thank the Francis Crick Institute Peptide Chemistry Facility for synthesis of fluorescent peptides for GINS labelling. This work was supported by the Francis Crick Institute, which receives core funding from Cancer Research UK (FC001221), the UK Medical Research Council (FC001221), and the Wellcome Trust (FC001221). G.C. was supported by a PhD Fellowship from Boehringer Ingelheim Fonds. K.N. and M.S. acknowledge the support from the Wellcome Trust (107935/Z/15/Z) and CRUK (26747).

MS Word template for this pre-print is from https://github.com/finkelsteinlab/BioRxiv-Template/blob/master/README.md.

## Author Contributions

M.S. purified labelled *Xenopus* cohesin complexes and performed experiments to test for recombinant cohesin loading in bulk. D.G. performed single-molecule experiments with Fen1-mKikGR comparing undepleted and cohesin depleted extracts. S.X. cloned and expressed *Xenopus* GINS. G.C. performed all other experiments. M.S and H.Y. supervised the research. G.C. and H.Y. wrote the manuscript with input from all authors.

## Materials & Correspondence

Correspondence should be addressed to M.S. or H.Y.

