## Supplementary Information for "Sister chromatid cohesion establishment during DNA replication termination"

*Supplementary Discussion*

*Number of cohesin complexes remaining at termination sites*

The number of cohesin complexes providing cohesion at termination sites is not evident from our experiments. We observe multiple cohesin rings remaining at positions where replication forks converge (for example, see Fig. 2a) but the presence of multiple cohesins is likely exaggerated as multiple cohesins are more likely to survive photobleaching. In instances where single cohesin rings are present after replication completion, it is not possible to exclude if unlabelled recombinant or endogenous cohesins are present.

*Interaction between tethered sister DNAs*

During sister DNA collapse experiments the time that both sister DNAs remain together before separating varies (Fig. 3b), even in cohesin-depleted extracts (Fig. 3d). Collapsed sister DNAs might not dissociate from one another immediately in cohesin-depleted extracts because of residual levels of endogenous cohesin. An alternative explanation is that there is cohesin-independent tethering of DNAs through binding of other proteins present in extracts.

Furthermore, we observe that when cohesin is present in extracts, there is a wide distribution of lifetimes that sister DNAs remain interacting. Sister DNAs might come apart because of the linear nature of the DNA used for our experiments, meaning that DNA can slide out of the cohesin ring (see Discussion). If collapsed DNA strands are resected sister DNAs could slide out of cohesin rings more easily. There is some evidence of DNA resection in our experiments – for example in the right-hand kymogram in Fig. 3b, fluorescent signal from collapsed DNA disappears more quickly that signal from the stretched DNA.

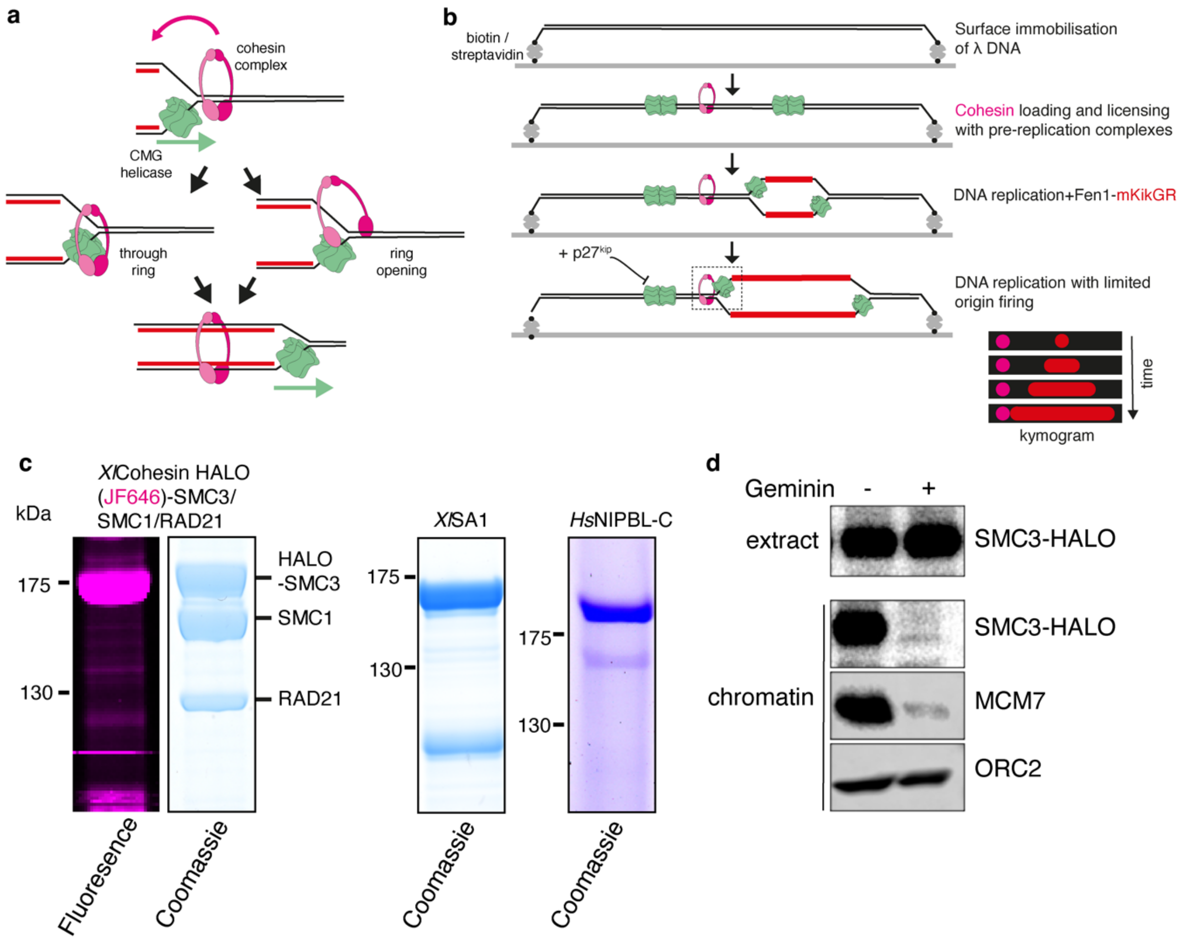


**Supplementary Figure 1. Characterisation of labelled Xenopus cohesin complexes**

**a**, Diagram showing the expected transfer of cohesin rings behind the replication fork, alongside possible models explaining how cohesin is transferred. **b**, Schematic of single-molecule replication assay, where Fen1-mKikGR is used to visualise replication fork collision with pre-loaded cohesin. **c**, Coomassie stained SDS-PAGE gels showing cohesin trimer (HALO(JF646)-SMC3/SMC1/RAD21), *Xl*SA1 and *Hs*NIPBL$^{c}$. **d**, Western blot showing Geminin sensitivity of JF646-cohesin loading onto chromatin in *Xenopus* egg extracts.

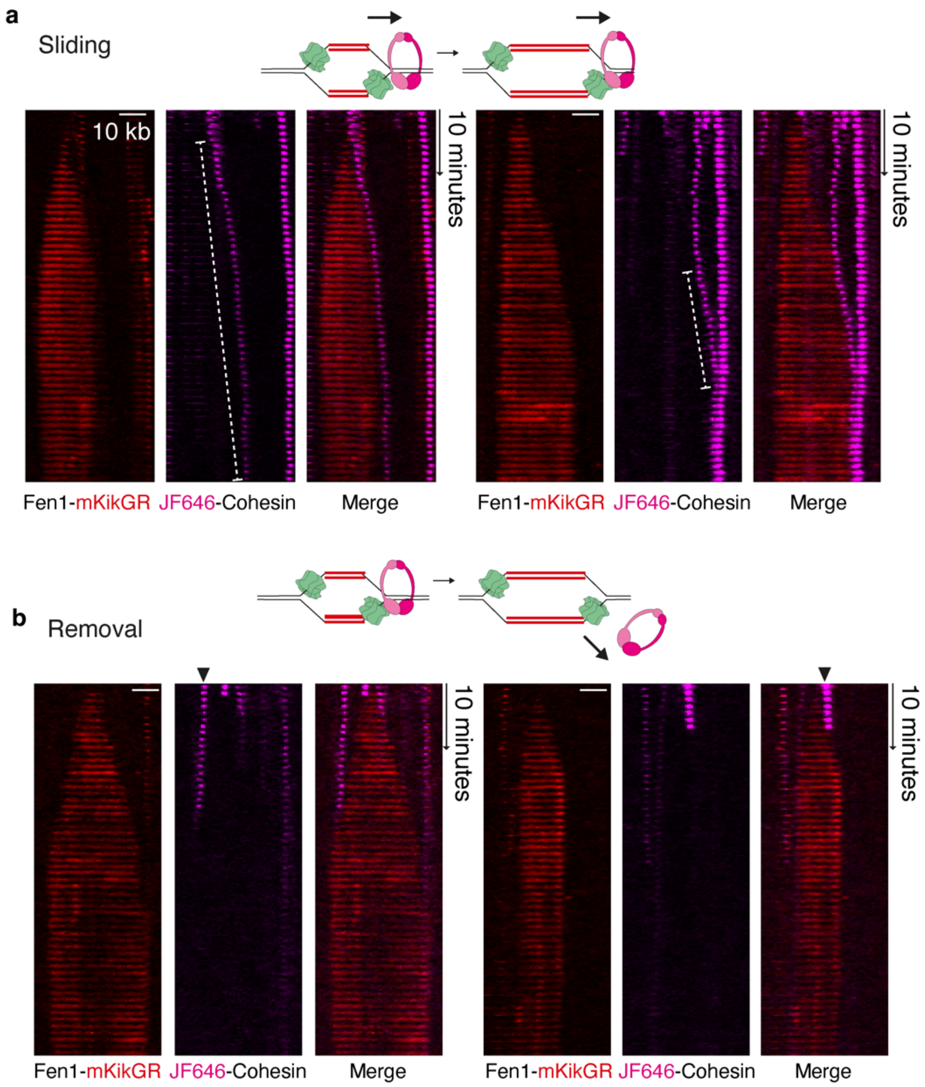


**Supplementary Figure 2. Cohesin is pushed ahead of replication forks in *Xenopus* egg extracts**
**a**, Kymograms showing real-time TIRF imaging of surface tethered λ DNA during replication from single origins in *Xenopus* egg extracts. In these examples JF646-cohesin (magenta) slides ahead of replication forks labelled with Fen1-mKikGR (red). **b**, Kymograms showing cohesin removal during replication from single origins.

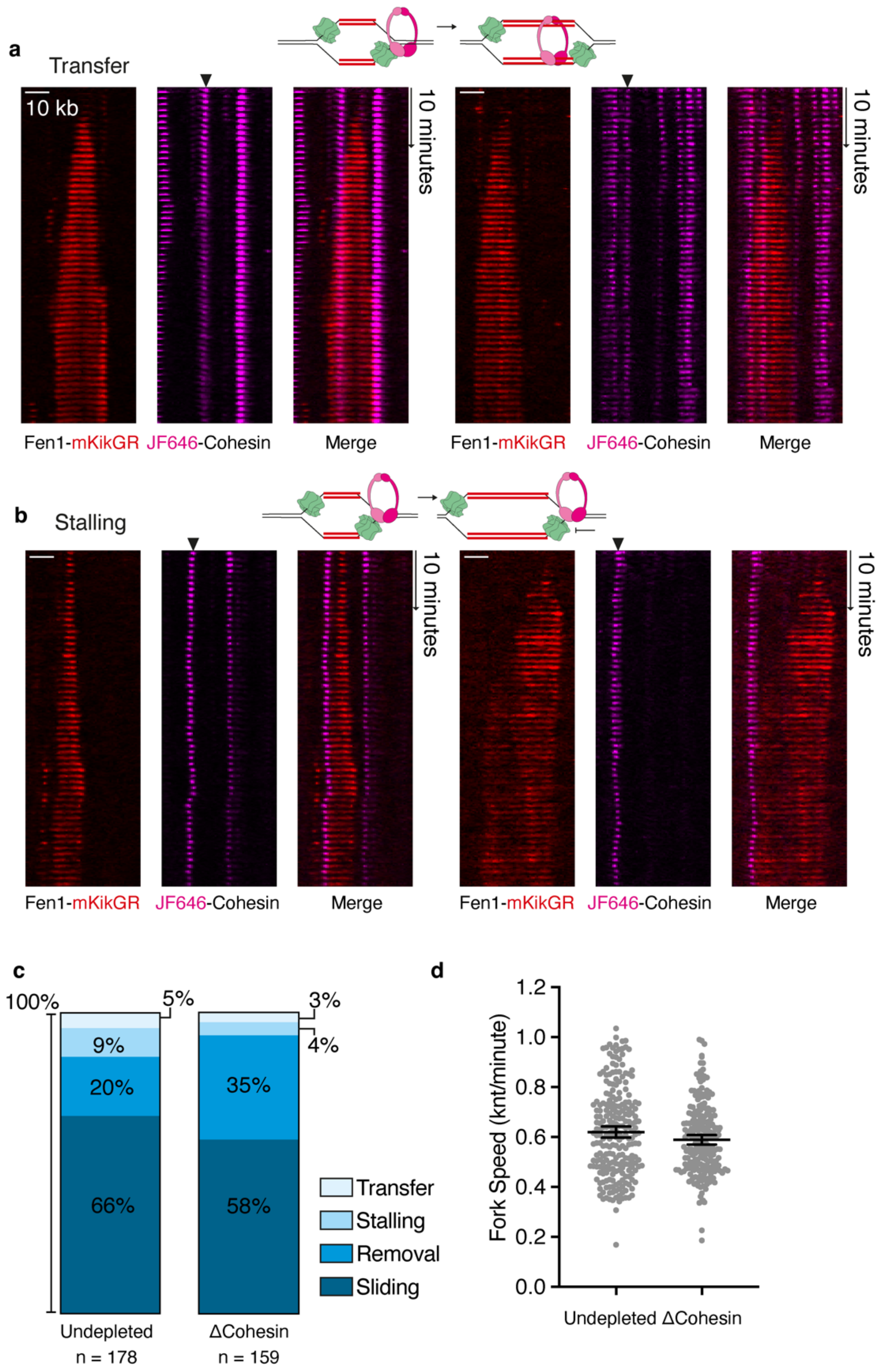


**Supplementary Figure 3. Replication fork stalling and cohesin transfer visualised during replication from single origins** Representative kymograms of **a,** cohesin transfer behind replication forks and **b,** replication fork stalling upon reaching cohesin. **c,** Comparison of primary cohesin fate upon collision by replication forks in normal *Xenopus* egg extracts compared to Δcohesin extracts. **d,** Quantification of fork speeds with Fen1-mKikGR in undepleted (0.62 kb/minute, n=237) and cohesin-depleted (0.59 kb/minute, n=220) extracts. Mean shown with 95% CI.

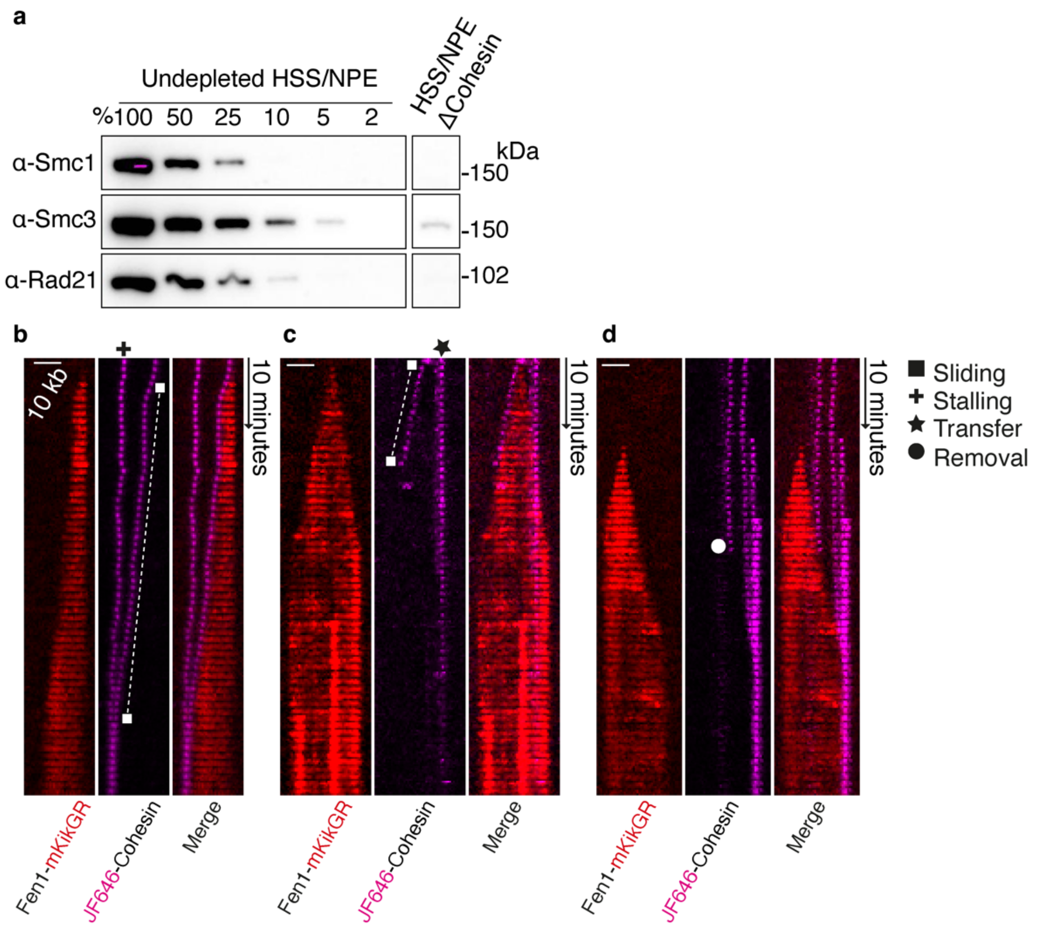


**Supplementary Figure 4. Cohesin depletion from Xenopus egg extracts**
**a**, Western blot showing >95% removal of Smc1, Smc3 and Rad21 subunits from replication extract after immunodepletion. **b-d**, Representative kymograms showing replication fork collision with pre-loaded cohesin in cohesin-depleted replication extracts. Origin firing is limited by addition of p27$^{kip}$.

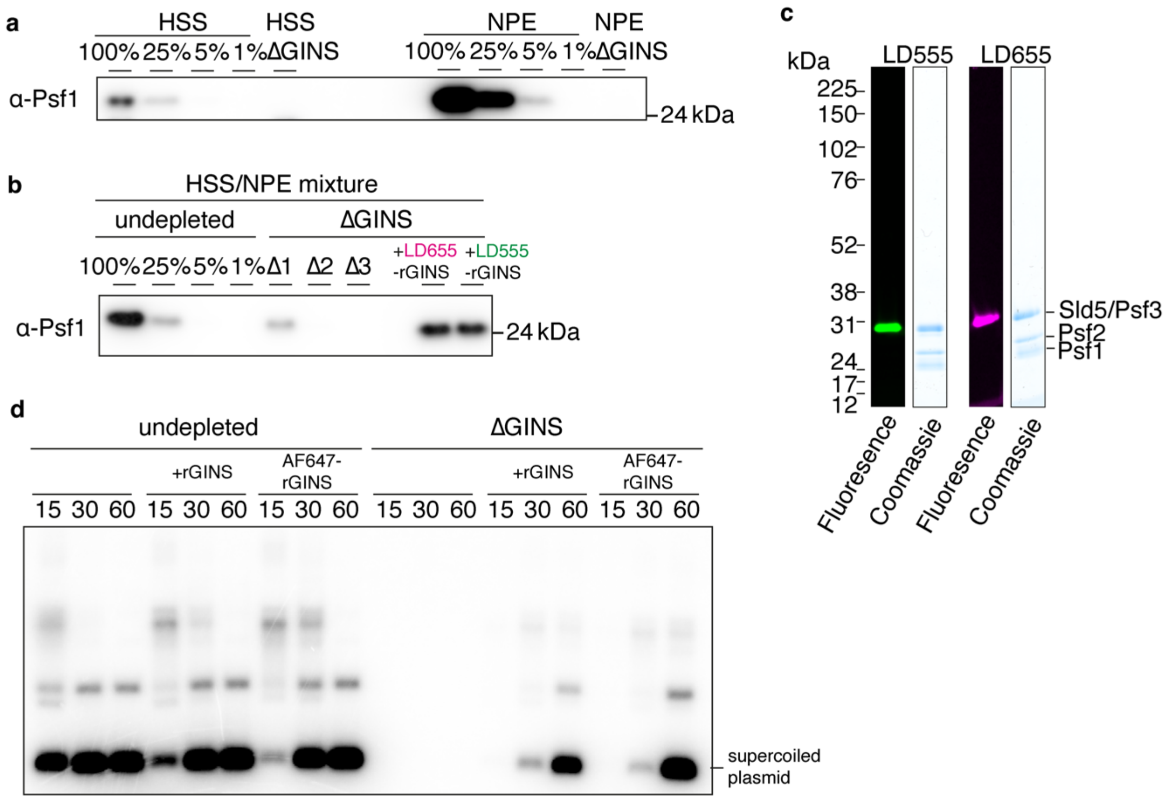


**Supplementary Figure 5. Establishing DNA replication in *Xenopus* egg extracts with labelled GINS**
**a**, Western blot showing GINS immunodepletion from HSS and NPE extracts separately (these extracts are used for bulk replication assays in d). **b**, Western blot showing GINS immunodepletion from a mixture of HSS and NPE (these extracts are used for single-molecule replication assays). **c**, SDS-PAGE gel coomassie stained / fluorescence scanned showing LD555- (green) and LD655- (magenta) labelled GINS complexes. **d**, Plasmid DNA replication in undepleted and GINS-depleted extracts with addition of recombinant GINS complexes. Plasmid DNA containing $^{32}$P-dATP was separated on a native agarose gel before visualisation by autoradiography.

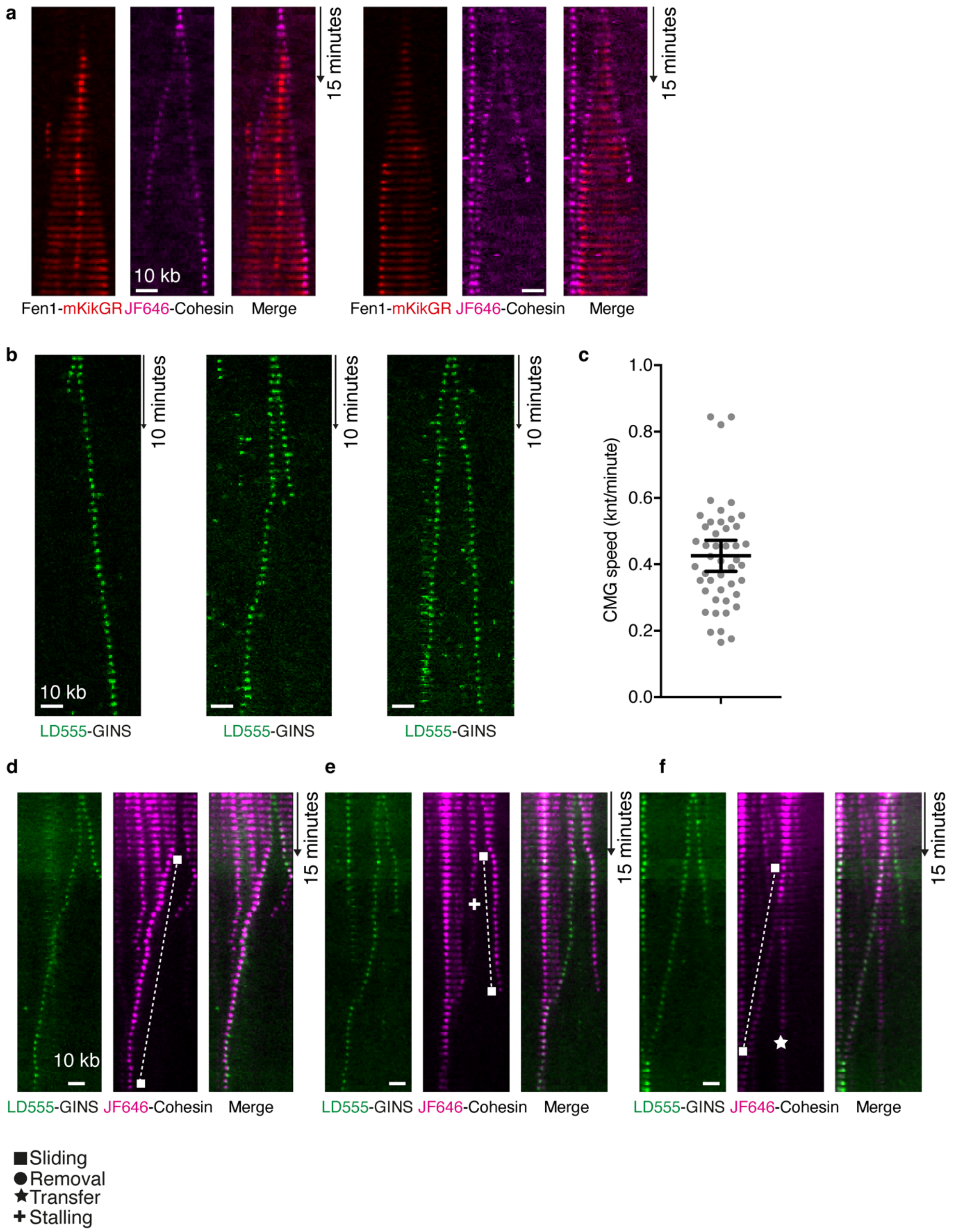


**Supplementary Figure 6. DNA replication with individual labelled replisomes**
**a**, Kymograms showing limited origin firing with replisomes containing LD655-GINS (magenta) and nascent DNA labelled with Fen1-mKikGR (red). **b**, Kymograms showing limited origin firing with replisomes containing LD555-GINS (green). **c**, Replisome speeds with LD555-GINS (n=46, mean=0.426 kb/minute, shown with 95% CI). **d**-**f**, Supplement to Fig. 2 with further examples of LD555-GINS / JF646-cohesin collisions after limited origin firing.

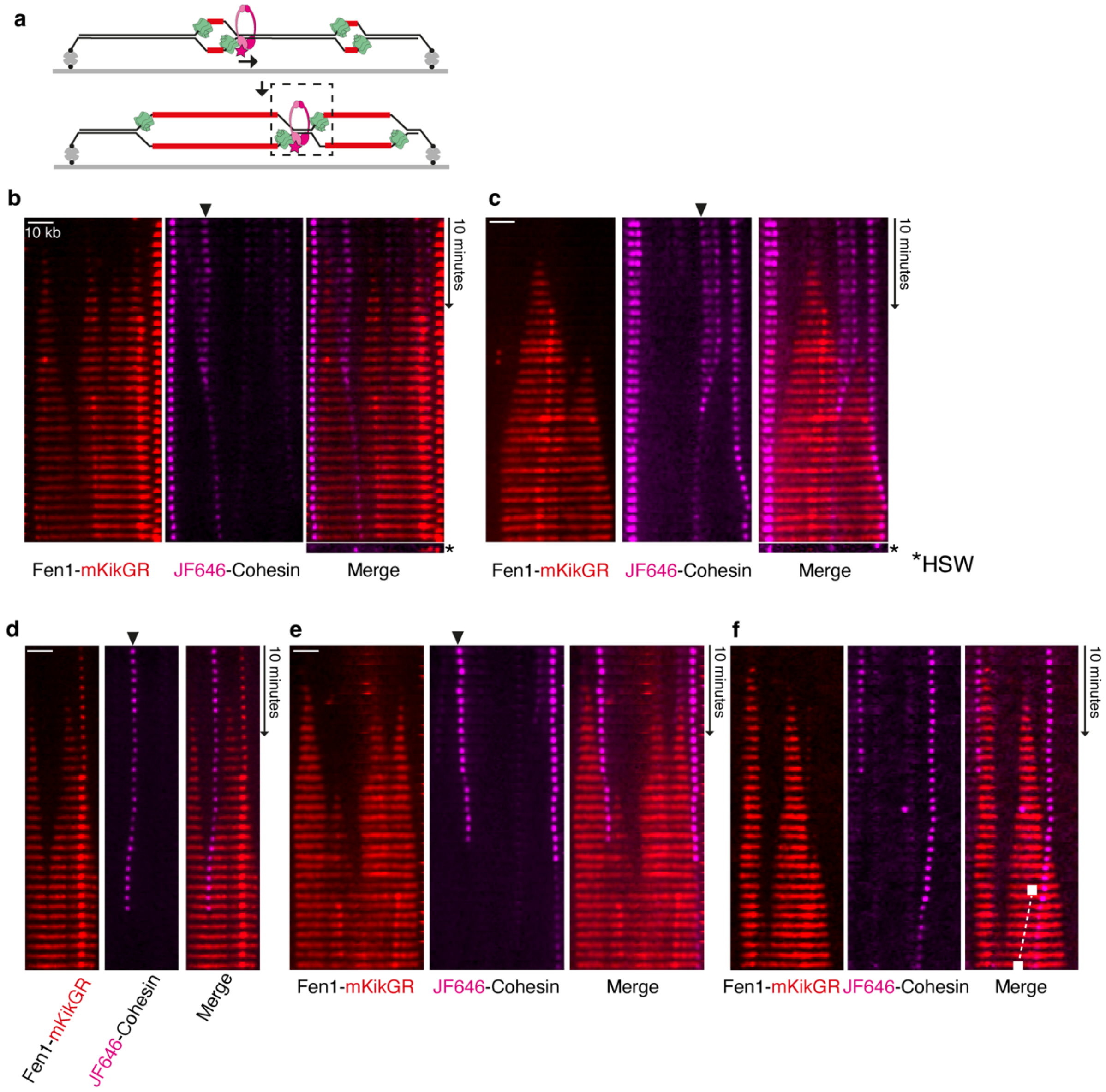


**Supplementary Figure 7. Cohesin fates at positions where replication forks converge**
**a**, Schematic showing DNA replication from multiple origins on tethered DNAs. Replication forks are marked with Fen1-mKikGR. Cohesin pushed ahead of a replication fork is visualised when meeting a converging replication fork. **b**-**f,** Supplement to Fig. 3, additional examples of JF646-cohesin fate at positions of converging Fen1-mKikGR labelled replication forks.

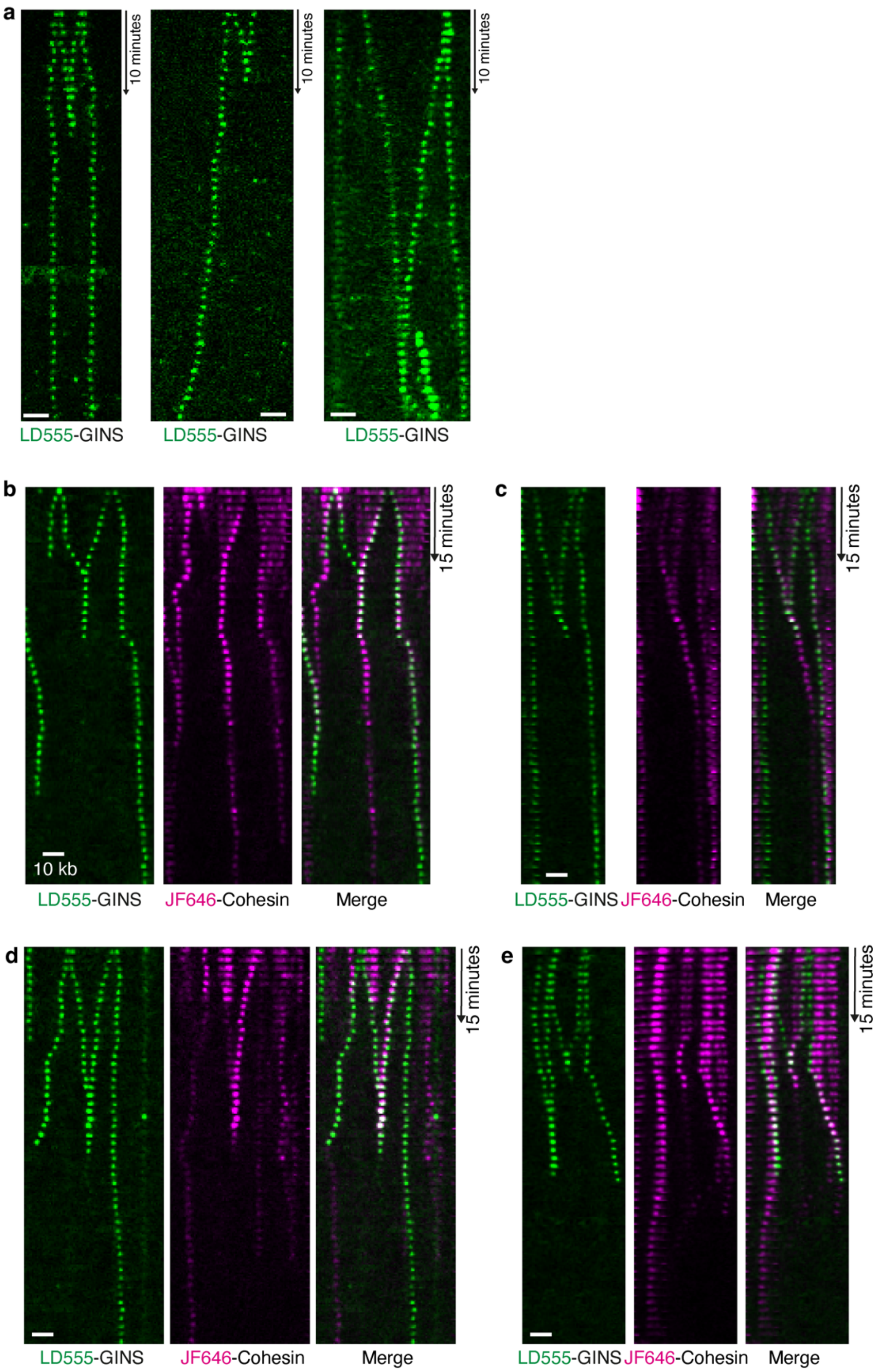


**Supplementary Figure 8. During DNA replication termination replisomes are removed and cohesin remains**
**a**, Kymograms showing DNA replication with LD555-GINS after firing from multiple origins. **b**-**e**, Additional representative examples of replisomes (LD555-GINS) pushing JF646-cohesin to sites of replication termination. In **b** cohesin remains, in **c** cohesin moves from the site of replisome disassembly. In **d-e** cohesin is removed.

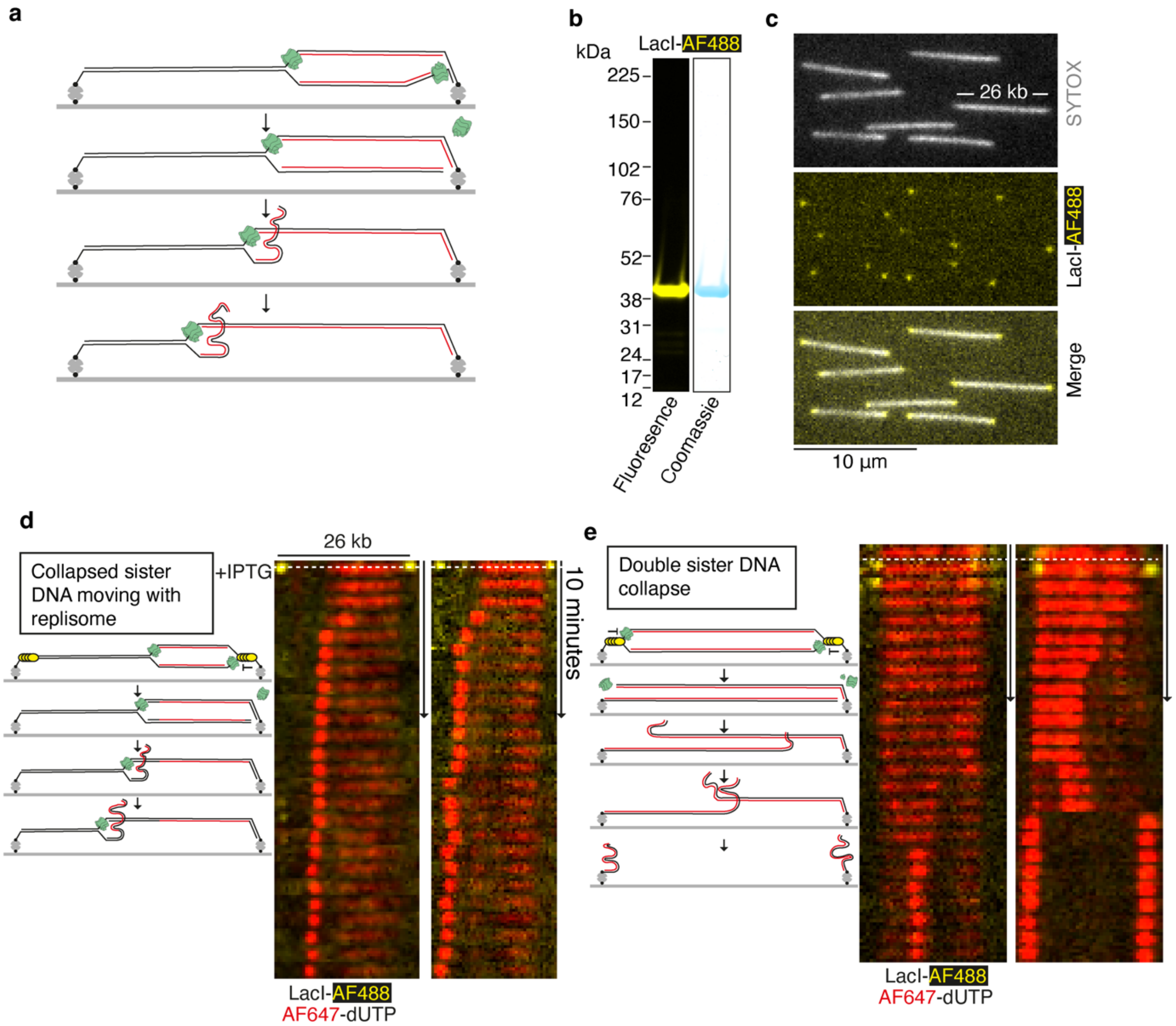


**Supplementary Figure 9. DNA templates to study sister DNA collapse after replication**
**a,** Schematic showing a newly replicated sister DNA being liberated from surface-tethered DNA. The collapsed sister DNA localises with the leftwards moving replisome. **b**, SDS-PAGE gel showing purified LacI-AF488. **c**, TIRF images of SYTOX Orange stained DNA bound by LacI-AF488. **d**, Kymogram examples of newly replicated sister DNAs collapsing and colocalising with the replisome. **e**, Examples where both newly replicated sister DNAs collapse to the same position on surface-tethered DNA. In the right-hand example, the two new sister DNAs completely separate.

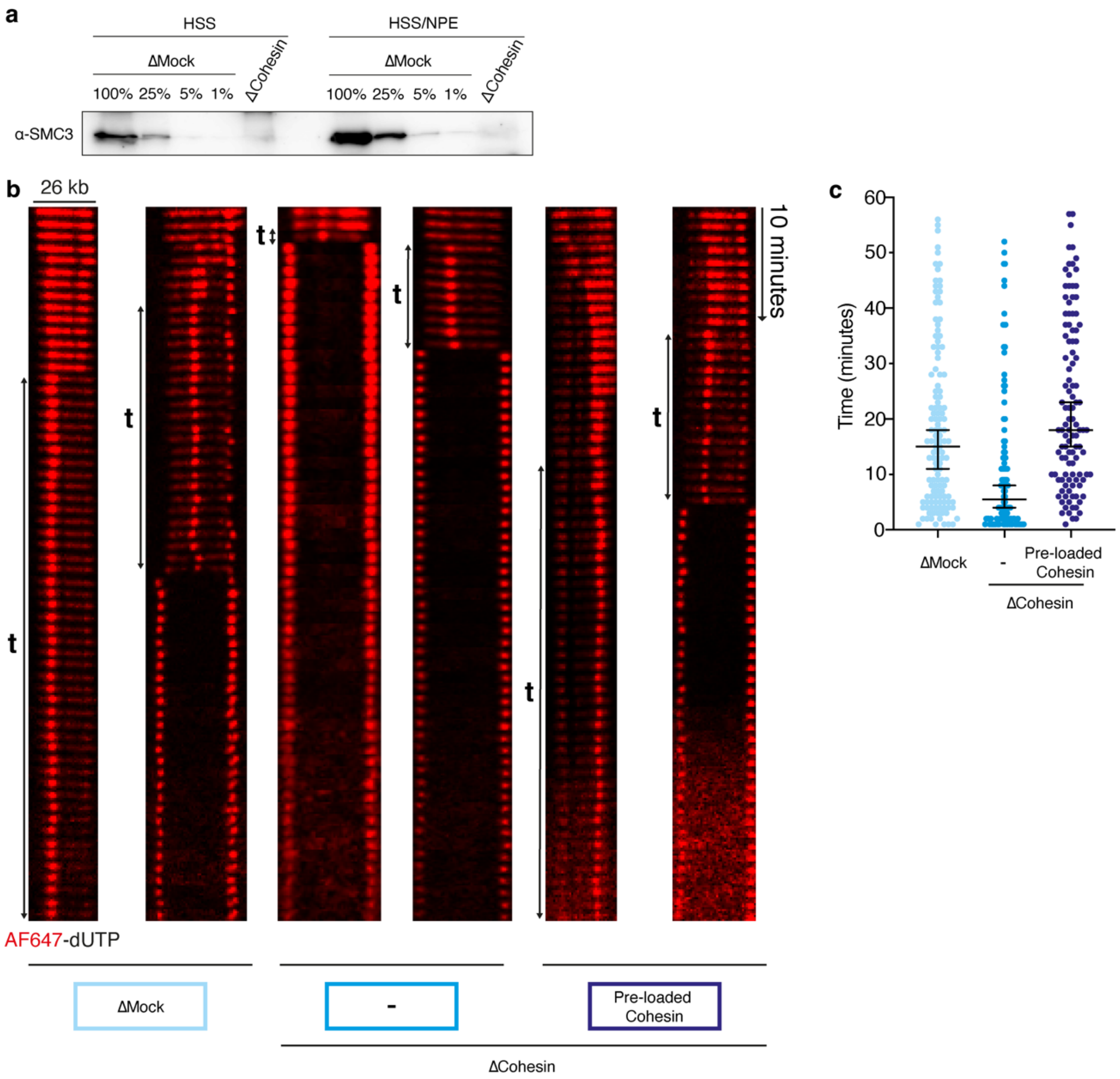


**Supplementary Figure 10. Immunodepletion shows cohesin dependent interaction between newly replicated sister DNAs**

**a**, Western blot showing mock-depleted and cohesin-depleted extracts used for DNA strand collapse experiments. HSS was used for DNA licensing and HSS/NPE mixtures were used for replication. **b**, Representative kymograms of double sister DNA collapse events where the time collapsed sister DNAs remain together is indicated. The conditions used for each example are indicated. **c**, Individual data points showing time that collapsed DNA strands remain together for mock-depleted (n=127), cohesin-depleted (n=106) and rescue (n=110) extracts. Data are shown mean with 95% CI.

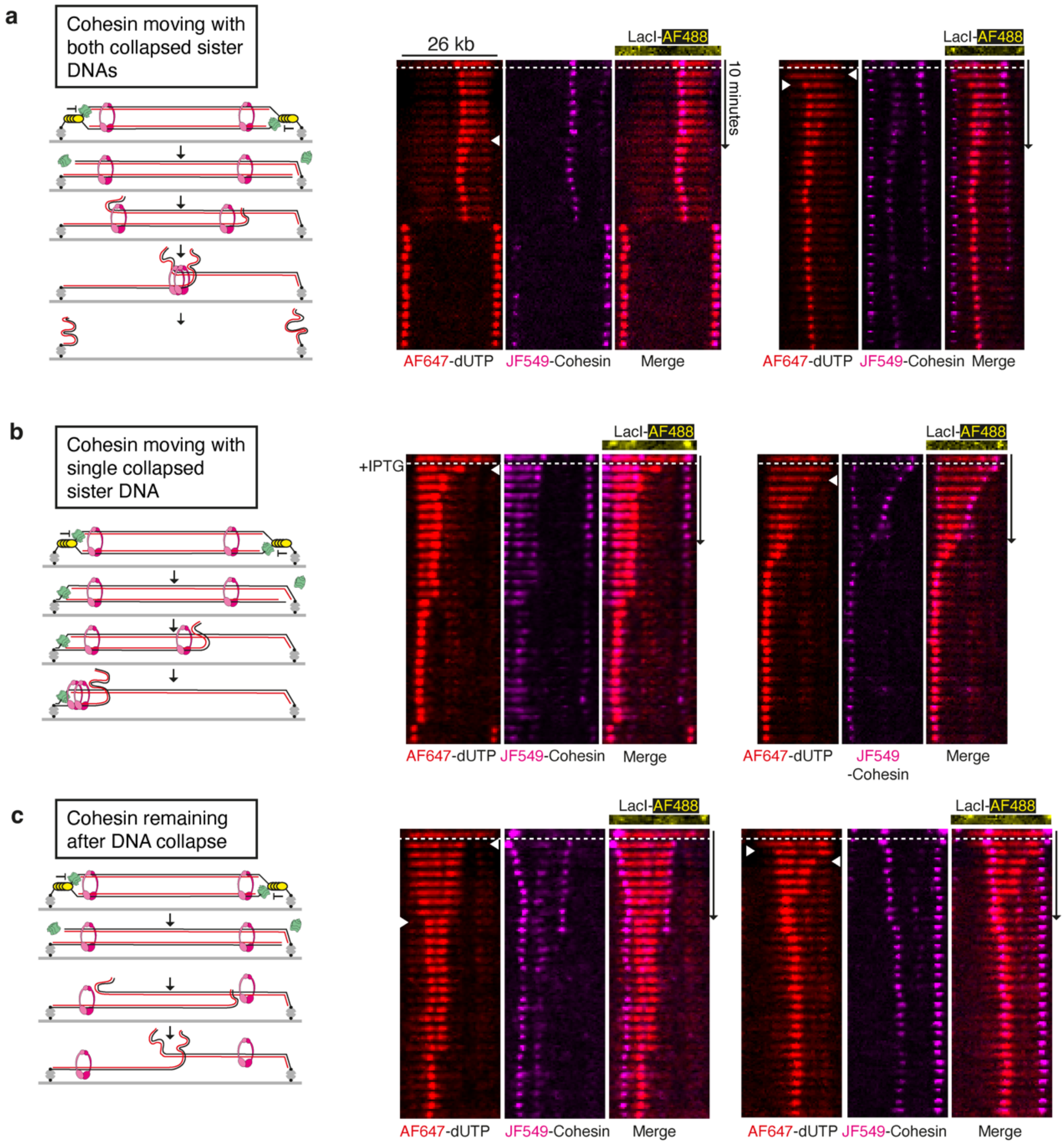


**Supplementary Figure 11. Cohesin fates after sister DNA collapse**

**a**, Kymogram examples showing cohesin moving with both collapsed DNA strands, as seen in Fig. 4b. **b**, Kymogram examples where cohesin moves with a single collapsed DNA strand, as seen in Fig. 4c. **c,** Kymogram examples where cohesin remains on surface tethered DNA after newly replicated DNA strands collapse.
